# Role of Nutritional Status on Arsenic Toxicity in *Daphnia pulex:* A Transcriptomic Perspective on Individual and Interactive Effects

**DOI:** 10.64898/2026.08.11.744190

**Authors:** Emily R. DeTemple, Craig E. Jackson, Anthony Schultz, Thomas H. Hampton, Joseph R. Shaw, Priyanka Roy Chowdhury

## Abstract

Inorganic arsenic is a widespread environmental contaminant and known human carcinogen, yet the mechanisms by which nutritional status modulates arsenic toxicity remain poorly understood. Here, we investigated the main and interactive effects of environmentally relevant concentrations of arsenic, low food quantity, and low dietary phosphorus supply on genome-wide gene expression in aquatic grazer *Daphnia pulex.* Differential gene expression analysis identified a total of 1,213 differently expressed genes with interactions of arsenic x nutrient stressors accounting for approximately 70% of the transcriptomic response. Low phosphorus emerged as a dominant main effect stressor and it also had a profound impact on transcription as a co-stressor. The low phosphorus x arsenic interaction exhibited the greatest transcriptional impact (435 DE genes), revealing that phosphorus limitation rather than food quantity influences arsenic toxicity at the gene expression level. Gene ontology and Pathway Activation Analysis revealed that main effects elicited simple yet distinct functional responses, whereas arsenic x nutrient interactions induced complex pathway-level disruptions including cell signaling, detoxification metabolism, DNA repair mechanisms, and energy homeostasis. Further assessment of gene expression revealed that all arsenic x nutrient interactions are antagonistic supporting previous literature that found arsenic behaves antagonistically as a co-stressor. Our results provide mechanistic insight into how nutritional status modulates arsenic toxicity and highlights the importance of considering arsenic x nutrient co-stressor interactions.

## Introduction

Arsenic compounds are ubiquitous environmental contaminants that pose a significant environmental and public health threat, primarily through contaminating drinking water. Due to their release via natural processes and anthropogenic activities, inorganic arsenic compounds have remained a high-priority toxicant for decades (Ravenscroft et al., 2009). Classified as a Group 1 human carcinogen since 1980, it is associated with various diseases including cancer, diabetes, and heart disease (IARC, 1980). The Agency for Toxic Substances and Disease Registry (ATSDR) has ranked arsenic first or second on its Substance Priority List for over two decades (Agency for Toxic Substances and Disease Registry, 2026) and it also appears among the World Health Organization’s (WHO) top ten chemicals of major public health concern (WHO, 2022). Given that an estimated 220 million people are exposed to high levels of arsenic in contaminated groundwater (Podgorski & Berg, 2020), and the health of our freshwater ecosystems are intrinsically tied to this issue, furthering our understanding of arsenic-elicited effects remains of high priority.

Chronic arsenic exposure produces varied effects across taxa such as impairment of metamorphosis in amphibians (Davey et al., 2008), elevation of oxidative stress in zebrafish (Yang et al., 2007), disruption to acclimation responses in killifish (Hampton et al., 2018), and reductions in growth and reproduction in invertebrates (Bi et al., 2024; Lin et al., 2025). The toxicity of arsenic is strongly influenced by its metabolism resulting in variation of effects due to arsenic disrupting multiple cellular and biomolecular pathways (Dreval et al., 2018; Tam et al, 2020). Several proposed mechanisms for arsenic toxicity include, but are not limited to, alterations in DNA methylation, disruptions in DNA repair mechanisms, increase in oxidative stress, and promotion of cell proliferation and apoptosis (Hughes 2002, Tam et al., 2020, Byeon et al., 2024, Bi et al., 2024). However, laboratory studies typically examine stressors like arsenic in isolation, which ultimately does not reflect the reality that organisms face multiple stressors in their respective environments (Folt et al., 1999; Crain et al., 2008; Piggott et al., 2015; Côté et al., 2016).

Among the many common environmental factors that modulate toxicant response, nutrient availability is particularly influential (Hennig et al., 2012). Several studies have reported that nutrient availability can play a significant role influencing how an organism responds to a given stressor. Copper tolerant and sensitive strains of *Scenedesmus acutus* experienced less inhibition of photosynthesis during copper exposure with increased levels of phosphorus(Twiss & Nalewajko, 1992). Similarly, Kaamoush and El-Agawany (2026) found that phosphorus limitation exacerbated the toxic effects of zinc and copper on the microalga *Dunaliella tertiolecta*, resulting in reduced growth and chlorophyll content. The essential nutrient phosphorus is of particular interest because arsenate [As(V)] is chemically analogous to phosphate, allowing arsenic to compete for phosphorus transport and binding sites through competition for transport pathways and cellular binding sites (Meharg and Hartley-Whitaker, 2002;Villa-Bellosta & Sorribas, 2010). Recent studies have also found a significant interaction between arsenic and dietary phosphorus, where *Daphnia* exhibited higher suppression of growth and reproduction under low phosphorus conditions than under optimal phosphorus conditions, when exposed to chronic arsenic (Schultz et al., 2024; Ayowemi et al., 2020). However, relatively little is known about how variation in nutrient availability influences arsenic toxicity in aquatic organisms at the molecular level. Such inquiries are especially relevant for freshwater ecosystems, that are vulnerable to sudden changes in both nutrient loading and arsenic accumulation (ATSDR, 1998; Dodds 2002).

Identifying interactions and characterizing co-stressor interactive effects is still considered one of the most pressing questions in ecology and conservation (Breitburg et al., 1998; Crain et al., 2008; Altschuler et al., 2015) with an increased emphasis on understanding how interactions influence the development of arsenic toxicity effects (Cheng et al., 2024). Distinguishing the interaction type of two co-stressors is a multifaceted problem within itself. The additive framework is a widely applicable model to assess co-stressor interactions (Folt et al., 1999). This model states that a co-stressor interaction can be synergistic (e.g. combined effect is greater than the sum of the effects) or antagonistic (e.g. combined effect is less than the sum of the effects).

Depending on the parameter of interest, there are multiple ways to interpret and use the defined terms (see Piggott et al., 2015; and Côté et al., 2016); however, deviations from additivity remain at the forefront of identifying co-stressor interactions (Folt et al., 1999). It is worth noting that many other variables can and may influence co-stressor interactions such as variability in toxicokinetic (e.g. stressor characteristics, intensity, frequency) and toxicodynamic (e.g. mechanism leading to effect produced) processes. Considering these variables can make predicting and managing co-stressor interactions an insurmountable task; increased research efforts have gone into assessing and characterizing general patterns with the most common environmental co-stressors (Crain et al., 2008; Piggott et al., 2015; Côté et al., 2016). Given arsenic’s prevalence in the environment, it’s important to continue dissecting such environmental contaminant co-stressor interactions, especially the potential rise of arsenic x nutrient interactions.

A growing body of evidence suggests arsenic most often act antagonistically with other stressors. Crain and co-authors (2008) compiled over 200 ecological studies to determine whether common co-stressor interactions (e.g. salinity, UV, temperature, toxicants, and nutrients) showcased predictive patterns. Collective data from the meta-analysis reported that toxicants, like arsenic, interacting with nutrients are generally antagonistic consistently produced opposing effects (Crain et al., 2008). Similarly, arsenic x selenium interactions are described to behave antagonistically causing cell apoptosis in human leukemia cell lines as well as inhibiting selenium uptake during gastrointestinal absorption (Ali et al., 2020). Also, multiple studies assessing arsenic-salinity interactions in killifish have reported antagonistic interactive effects interfering with essential processes related to salinity acclimation at the multiple levels of biological organization (Shaw et al., 2014; Hampton et al., 2019). In a study assessing a natural population of *Daphnia magna*, arsenic x copper interaction exhibited a reduced effect on reproduction than either single stressor response (Asselman et al., 2019).

Impacts from interactive effects on organismal fitness are typically measured for growth, reproduction, and survival (Sales et al., 2016; Lind et al., 2018; Awoyemi et al., 2020; Schultz et al., 2024). However, transcriptional responses precede the observable physiological changes making gene expression an early indicator of stressor and interactive effects (Shaw et al., 2007; Quirós et al., 2007). Gene expression data has been used to assess binary metal interactions in *Daphnia* species (Vandenbrouck et al., 2009; Asselman et al., 2019) and calcium-predator presence on target genes in *Daphnia pulex* (Altshuler et al., 2015). Similarly, several studies have shown that organisms frequently adjust their transcriptional profiles in response to variations in nutrient availability altering expression of genes involved in nutrient acquisition, energy metabolism, protein synthesis, antioxidant defense, and stress response pathways (Jeyasingh et al., 2011; Xu et al., 2021). For example, nitrogen limitation triggers upregulation of genes involved in nitrogen uptake and stress-response pathways in photosynthetic organisms (Allen et al., 2008; Amtmann and Armengaud, 2009).

Similarly, dietary restrictions in *Drosophila melanogaster* alters genes associated with insulin signaling, energy metabolism and stress response (Partridge et al., 2005), while nutrient limitation in the freshwater zooplankton *Daphnia* induces transcriptional changes in pathways related to phosphorus metabolism, growth, and reproduction (Jeyasingh et al., 2011; Roy Chowdhury et al., 2014; Xu et al., 2021).

In this study, we tested the interactive effect of sublethal, environmentally relevant arsenic concentrations, low food quantity (i.e., dietary restriction), and low dietary phosphorus on gene expression patterns in the aquatic grazer *Daphnia pulex*. The freshwater cladoceran *D. pulex* provides an ideal model system to investigate these interactions because of its ecological importance and well-established use in genome expression profiling (Shaw et al., 2008; Kim et. al., 2015). The main effects of food quantity (food), dietary phosphorus, and arsenic toxicity on *Daphnia* fitness are well documented (Food quantity: Latta et al., 2011; Vighi et al., 2003; Phosphorus limitation: Jeyasingh & Weider, 2005; Jeyasingh et al., 2011; Roy Chowdhury et al., 2014; Arsenic exposure: Shaw et al., 2007; Theegala et al., 2006). Recent studies have also found a positive interaction between arsenic and dietary phosphorus limitation, where *Daphnia* exhibited higher suppression of growth and reproduction under low-phosphorus conditions when exposed to chronic arsenic in comparison to optimal phosphorus conditions (Schultz et al., 2024; Ayowemi et al., 2020). Furthermore, limited food supply also altered somatic arsenic accumulation in *Daphnia* (Miao et al., 2012; Schultz et al., 2024). Genome-wide expression studies in *Daphnia*, also indicated significant impacts on gene expression in these organisms when exposed to dietary restriction or environmentally relevant arsenic concentrations. For example, Asselman et al. (2019) showed that in a natural population of *Daphnia,* gene families associated with stress responses, such as cuticle proteins, respond transcriptionally to arsenic exposure.

Similarly, elevated gene expression levels in stress related pathways were also observed under food limitation in *D. pulex* (Becker et al., 2018). Limited phosphorous supply also triggered differential expression in ∼10% of the *D. pulex* genome impacting genes involved in phosphate transport, drug detoxification and heavy metal uptake and absorption pathways (Jeyasingh et al., 2011; Roy Chowdhury et al., 2014). Taken together, these studies highlight *Daphnia* as an ideal model for investigating interactions among arsenic exposure, food quantity, and dietary P-supply shape gene expression patterns.

The main objective of the current study is to measure the individual and combined effects of chronic exposure to an environmentally relevant concentration of arsenic, low food quantity, and low dietary phosphorus on gene expression and biomolecular pathways in *D. pulex*. The present study is strengthened by a companion study that followed the same experimental design, yet focused on life-history/fitness parameters, i.e. reproduction (Schultz et. al, 2024). The authors report: (1) phosphorus limitation altered biological processes related to growth, (2) arsenic impacted molecular mechanisms critical to reproduction, and (3) low phosphorus x arsenic interactions allowed for earlier maturation in comparison to the low food x arsenic interactions.

Based on previous studies (Roy Chowdhury et al., 2014; Shaw et al., 2014; Hampton et al., 2018), we hypothesize that *Daphnia* will exhibit antagonistic gene expression patterns in the presence of arsenic x nutrient costressors. We predict interactive effects between arsenic x nutrient costressors will produce transcriptional responses conditional on the nutrient stressor. Further, given that arsenic competes with phosphorus for uptake (Miao et al., 2012), we predict that low phosphorus x arsenic interactive effects will identify gene pathways that commonly regulate arsenic metabolism and are conditional on the presence of phosphorus limitation. Quantifying stressor and stressor interactions effects on gene expression that can be interpreted in light of organismal fitness is a major strength of the present study and helps strengthen the mechanistic underpinnings of the arsenic x nutrient interaction.

## Methods

### Algal cultures for experimental feeding

Based on the specific food treatment, *Daphnia* was fed *Scenedesmus acutus* cultured in semi-continuous chemostats in either the optimal phosphorus medium (hereafter termed: HP; dilution rate of 1 d^−1^; Kilham et al., 1998) or the low phosphorus medium(hereafter termed: LP; dilution rate of 0.15 d^−1^; Kilham et al., 1998). These conditions produced algal diets with phosphorus concentrations of 50µmol L^−1^ and 5µmol L^−1^ (Roy Chowdhury et al., 2014). After chemostats reached a stable state (∼10 days), the algal outflow was collected to determine C-content spectrophotometrically before all experiments (Schultz et al., 2024).

### Daphnia cultures and maintenance

The *Daphnia pulex* clone used in this study originated from a pond in northwestern Iowa (Weider et al., 2004) and was maintained in the laboratory for 10 years at 20 ± 1°C, and 16:8 light: dark cycle in COMBO media (Kilham et al. 1998). All cultures were fed ∼3 mg CL^−1^ of optimal phosphorus algae. To obtain experimental organisms, 50 –60 gravid females were isolated from the above cultures and fed HP algae at 1mgC L^−1^ d^−1^. Less than 24 hour old neonates obtained from gravid females were used for all subsequent experiments.

### Arsenic exposure and RNA extraction

For samples used in our RNASeq experiment, > 24 hour old neonates from experimental cultures were divided into eight different treatment conditions following the experimental design (see Schultz et al. 2024). The sublethal concentration of arsenic (0.132 mg L^−1^) utilized in this study was determined from acute and chronic toxicity testing conducted in Schultz et al., 2024. There were two batches – one with media containing arsenic and one in which no arsenic was added. Arsenic solutions were prepared using reagent-grade sodium arsenate (Sigma-Aldrich; Cat. No. A6756) dissolved in COMBO media. Arsenic concentrations in media were analyzed prior to exposures in the Dartmouth Trace Metal Lab using inductively coupled plasma-mass spectrometry (ICP-MS; Agilent model 7900). Within each batch, there were two food concentrations - 3 mgC L^−1^ (well above saturating food concentration; Lampert 1977) and 0.1 mgC L^−1^ (below incipient food level). Within each food concentration, two levels of dietary phosphorus, 50 µmol L^−1^ and 5 µmol L^−1^, were introduced. Each treatment was replicated 3 times generating 24 transcriptomic profiles (see Schultz et al., 2024 for experimental design). Ten individuals were exposed to their respective treatment for 10 days. After 10 days, total RNA was extracted using the Qiagen RNeasy MiniKit (CAT No. 74104) and QIAshredder (Cat No. 79654) following the manufacturer’s protocol.

RNA quality (260/280 ratio: 2.0 - 2.3) and quantity (2.2 - 27.2 ng/uL ) were checked using NanoDrop (Invitrogen, Carlsbad, CA, USA) and Qubit^TM^ 4 Fluorometer (Thermo Fisher, Cambridge, MA) respectively, before shipping to UNH Hubbard Center for Genome studies for sequencing (https://hcgs.unh.edu/). Following the manufacturer’s protocols, RNA was first reverse transcribed using the SuperScript Double-Stranded cDNA Synthesis Kit (Thermo Fisher, Cambridge, MA). RNA-seq libraries were then constructed following a polyA selection approach using Illumina compatible Nextera DNA Flex Library Prep Kit (Cat. No. 20018705). Illumina-compatible adapters from the Nextera DNA Unique Dual Indexes (Cat. No. 20027213) were used to attach individual barcodes to all 24 libraries. Library size distribution was determined using a Bioanalyzer 2100 (Agilent Technologies, Santa Clara CA, USA) with DNA High-Sensitivity chips and reagents (Agilent Technologies, Santa Clara CA, USA). Illumina TruSeq SBS v4 reagent kit (300 cycles) was used to generate paired-end 250 bp reads using the Illumina HiSeq platform (Illumina, San Diego CA, USA).

### Sequencing Information

Initial quality check was done with *FastQC* (v0.11.6). Low quality reads were trimmed to a minimum Phred score of 30 using *Trimmomatic* (v 0.27) and adapters were removed with the following specifications (ILLUMINACLIP:TruSeq3-PE-2.fa:2:30:10:8 TRUE LEADING:20 TRAILING:20 MINLEN:40) (Bolger et al. 2014). The 24 *D. pulex* RNA samples, encompassing control and 8 treatment groups combinations, were sequenced in triplicate, generating a total of 25.4 million paired end reads with sequence length ranging from 40-251 base pairs. An average of 2,087,120 high-quality sequence reads per sample were obtained with a range from 430,896 – 3,558,805 reads for individual samples (Supplemental Table 1). One sample (Sample_9) indicated significantly lower input reads after trimming (27093) and overrepresented sequences indicating potential DNA/rRNA contamination resulting in its omission from all subsequent analysis (Supplemental Table 1). An average of 98.6% of the total reads (50.8 million) were used for downstream analysis. The transcriptomic index was created using reference transcripts from the National Center for Biotechnology Institute (NCBI) *Daphnia pulex* KAP4 GTF file (https://ftp.ncbi.nlm.nih.gov/genomes/all/annotation_releases/6669/100/GCF_021134715.1_ASM2113471v1/). The mapping based mode of *Salmon* (version 1.9.0) was used to quantify the transcripts applying default parameters (Patro et al., 2017). On average, 65.1% of reads per library mapped successfully with rates ranging from 30.6% - 79% across samples due to residual ribosomal RNA and mitochondrial RNA in libraries (Table S1).

### Preprocessing Data and Exploratory Analysis

Transcript-to-gene mapping was conducted from quantified transcripts and the NCBI *Daphnia pulex* KAP4 GTF annotation file using *GenomicFeatures R* package.

Transcript-level abundances were then imported and aggregated to gene-level counts using *tximport* following the recommendation of Soneson et al. (2016). Exploratory analysis was conducted in *R* (Robinson et al., 2010) to assess library size, count distribution, and clustering of raw and normalized data. Assessment of raw library size suggested that three of the 24 samples had fewer counts than 500,000. Further analysis of the log transformed normalized counts indicated that one sample remained significantly below median count (Fig. S1), and hierarchal clustering indicated a lack of clustering with any of the samples prompting its removal from the analysis as an outlier (Fig. S2). Principal component analysis was conducted on the treatment replicates after the removal of the outlier and clustering based on phosphorus treatment (Fig. S3).

### Differential Gene Expression Analysis

Differential gene expression analysis was conducted using the software package *edgeR* (Robinson et al., 2010). Genes were filtered using the “filterByExpr” function with the following minimum count thresholds (min.count = 10, total.min.count = 15), normalization factors were calculated for each library using the “calcNormFactors”, and data was normalized using counts per million (cpm) function. Log_2_-transformed expression values were fit to a negative binomial generalized linear model (glmFit) including all main effects (arsenic, low phosphorus, low food) and their subsequent binary interactions. Likelihood ratio tests as well as contrasts for each main effect and binary interaction term were conducted. The “decideTestsDGE” function was used to determine differentially expressed (DE) genes and adjusted for multiple testing using the false discovery rate (FDR) < 0.05 and log_2_fold change of 2. Threshold criteria were chosen in combination to assess statistically significant and biologically relevant genes.

Long non-coding protein genes were removed prior to annotation for downstream analyses (Table S2). Lists for annotated main effects and interactions terms can be found in the supplements (Tables S3-8). Venn diagrams were created to further explore DE genes that were shared between main effects and interaction terms and created using SRplot (Fig. 1)(Vandenbrouck et al., 2009; Tang et al., 2023).

**Figure 1:**
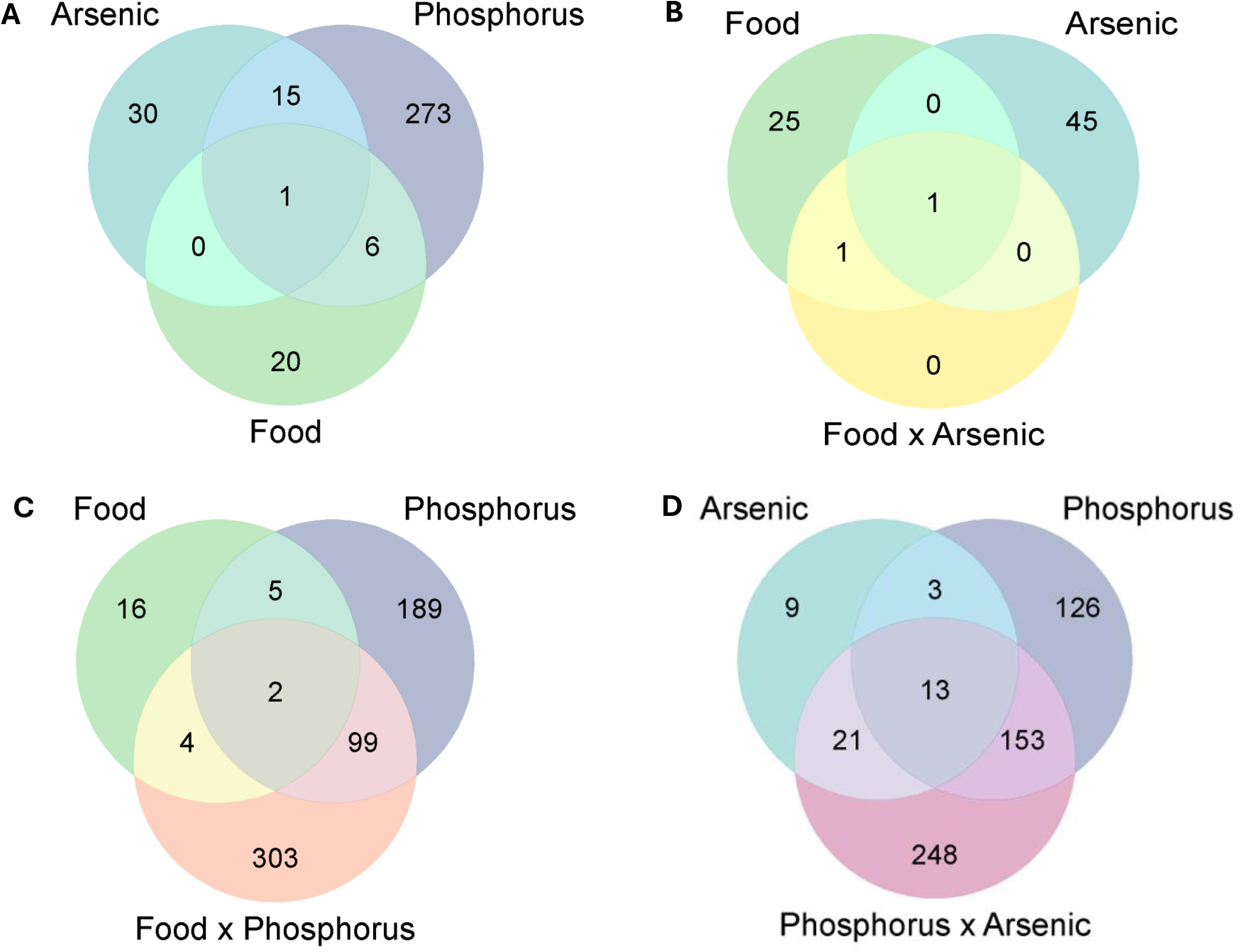
Differences in gene expression were analyzed by measuring quantile normalized log_2_expression values using a three-factor linear model that includes three variables (presence or absence of arsenic, low or optimal food quantity, low or optimal phosphorous concentrations in food) as well as all interactions. Due to complexity of interpreting interactions, this study focuses specifically on binary interactions. The descriptor “low” was removed for spacing purposes. Differentially expressed (DE) genes were defined by *p-*values < 0.05 and fold changes >2. Venn diagrams were constructed to visualize overlap in DE genes between main effects (A) and interaction terms with their constituent main effects (B-D). Total number of DE genes for the main effects (A) include arsenic (46 DE genes), low phosphorus (295 DE genes), and low food (27 DE genes). Total number of DE for each interaction term include food x arsenic (2 DE genes), food x phosphorus (408 DE genes), and phosphorus x arsenic (435 DE genes) with the increase in DE genes used to order the Venn diagrams from the lowest number of DE genes (B) to highest (D).

### Gene Ontology (GO) Functional Enrichment Analysis

Gene sets for main effects and interactions were assessed for functional enrichment using gProfiler’s Gene Ontology (GO) enrichment analysis tool. The goal was to evaluate which biological processes (BP), molecular functions (MF), and cellular components (CC) were impacted by main effects of arsenic x nutrient stressors and their binary interactions. The analysis was conducted by uploading unranked lists of differentially expressed NCBI Gene IDs for each main effect and interaction into the functional profiling g:GOSt tool on the g;Profiler web interface (parameters: organism = *Daphnia pulex* KAP4; advanced options: Benjamini-Hochberg FDR, Numeric ID: GENEID) (Kolberg et al., 2023). Bubble enrichment plots were created to highlight driver terms for each effect and were created using the SRplot web interface (Fig. 3 & 4) (Tang et al., 2023).

### Pathway Activation Analysis (PAA)

Pathway-level analysis provides a systems-level perspective on how gene expression changes translate into biological responses to environmental stressors. Unlike overrepresentation analysis (ORA), which relies on arbitrary significance thresholds and only considers differentially expressed genes, pathway activation analysis (PAA) leverages the full gene expression dataset to predict the directional activation or suppression of biological pathways, thereby improving sensitivity and biological interpretation. Consequently, PAA is particularly well suited for evaluating arsenic x nutrient interactions with relatively few differentially expressed genes, enabling the detection of coordinated pathway-level responses that would likely be overlooked by ORA. PAA utilizes a binomial test to determine if the proportion of induced/repressed genes in a pathway deviates from the expected 50:50 split (Hampton et al., 2018). In previous studies, this method assessed the systemic activation or suppression of biological pathways in the context of both individual and interacting co-stressors and provided critical insight into interpreting response severity and resultant observable effects on the model organism (Hampton et al., 2018). Pathways were deemed significant if more than 4 genes in a pathway were activated or repressed following threshold criteria of FDR < 0.05 & median log_2_fold change > 1. Tables exhibiting number of significant pathways for main effects can be found in the supplements (Table S9) and interactions found in the main text (Table 1).

**Table 1:**
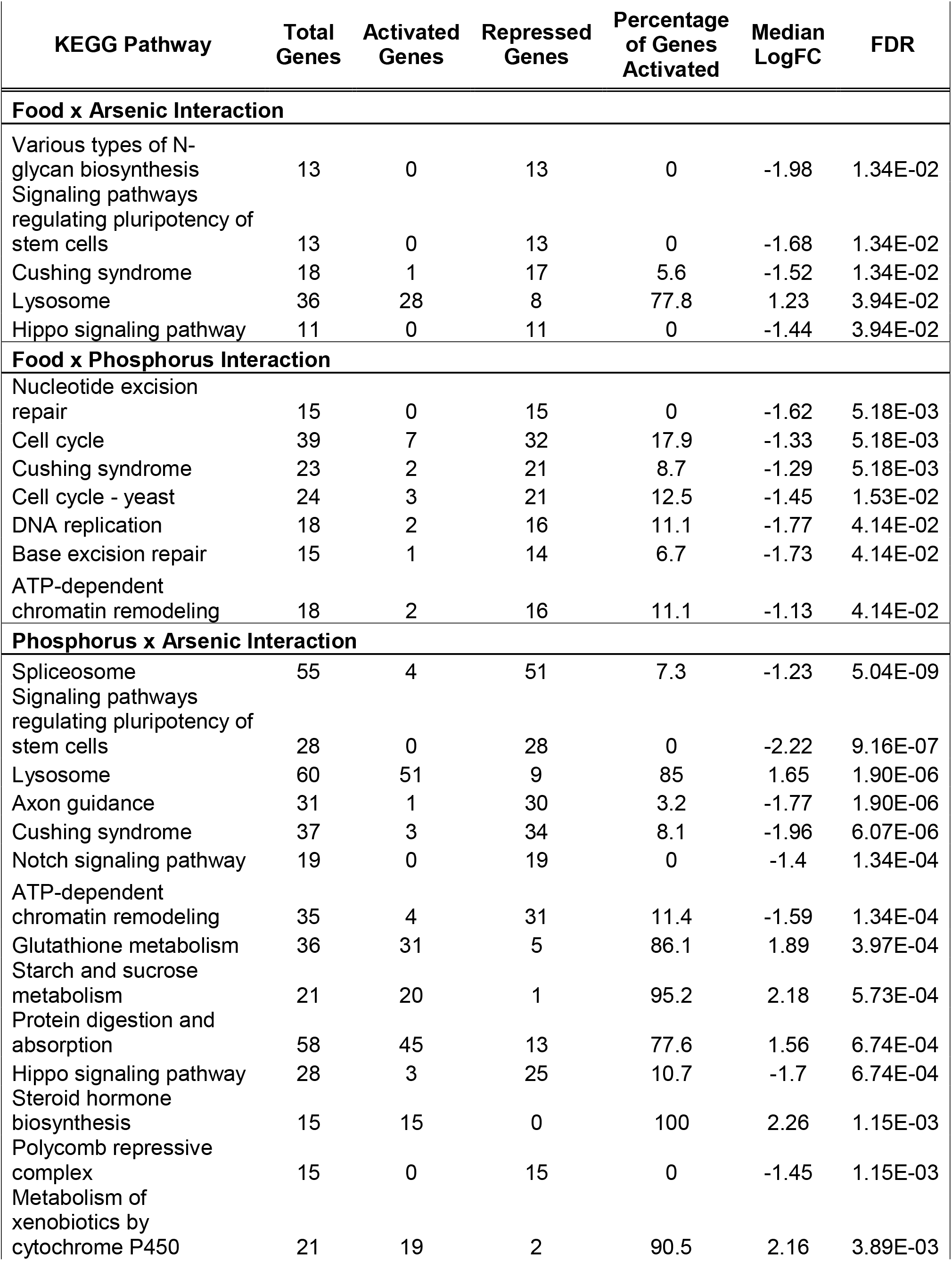
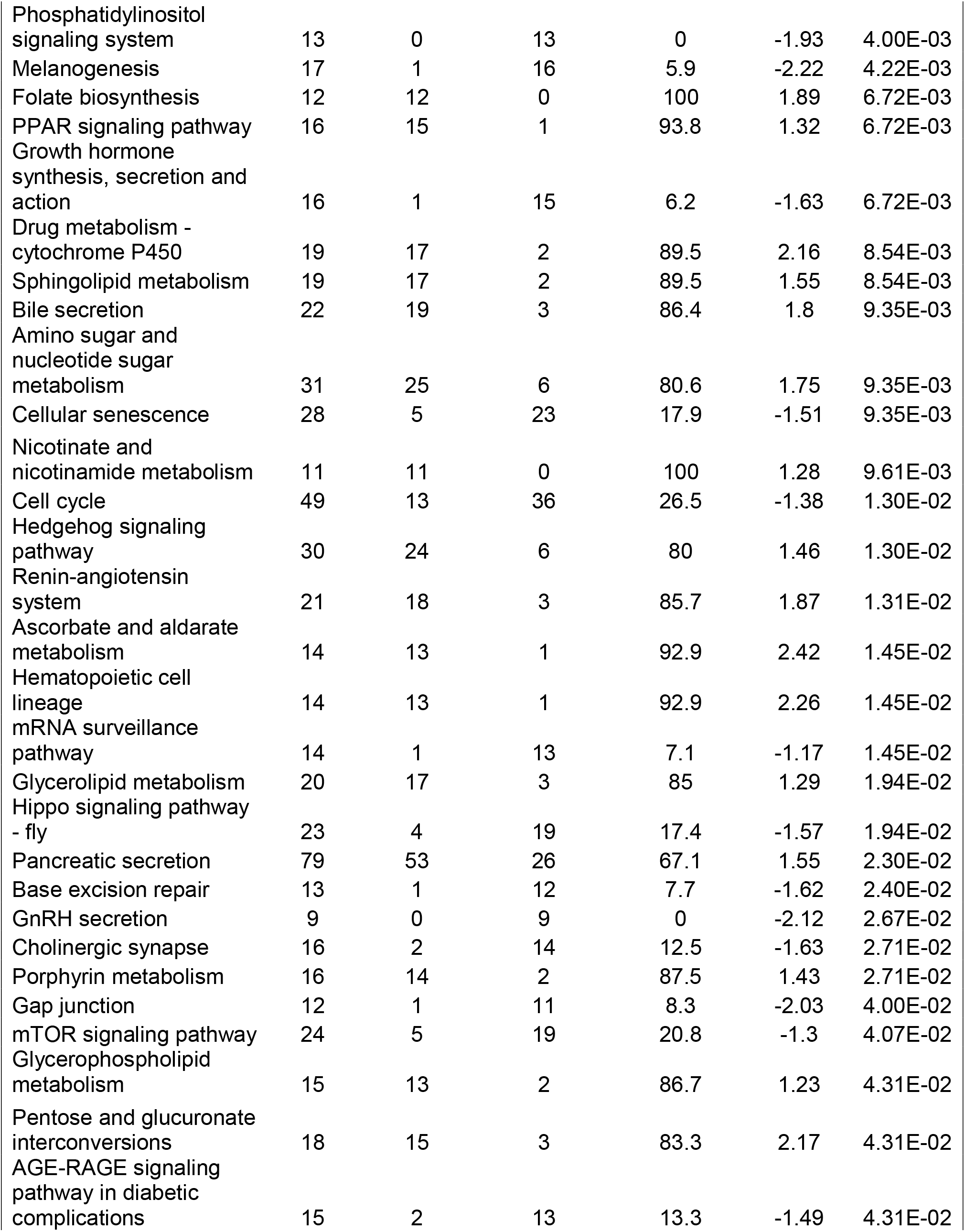
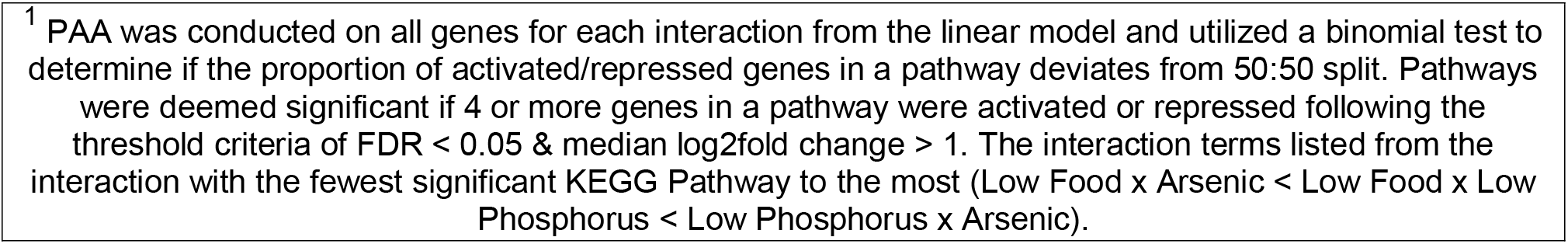
Pathway Activation Analysis Results for Binary Interactions^1^.

### Identification of Co-stressor Interaction Types via Gene Expression Patterns

Gene expression dynamics for each main effect and subsequent interactions were visualized to identify and characterize whether the *D. pulex* responses to arsenic x nutrient co-stressors were antagonistic or synergistic. From the full linear model, differential gene expression data for each main effect and subsequent co-stressor interaction were visualized to understand the magnitude and direction of each interaction (Fig.5, Fig. S4). Each subplot lists both main effects that comprise a given interaction from left to right on the x-axis. Subplots showcase DE gene expression dynamics for each main effect (Fig. 5A, B, D, & E; Fig S4A & 4B) and binary interaction (Fig. 5C & F; Fig. S4C) with each DE gene from the particular list of interest being represented by a red (up-regulated) or blue (down-regulated) line. We utilized an additive model to characterize all of the interactions; thus, the previously mentioned definitions for additivity, synergism, and antagonism were applied in the identification of interaction type.

## Results

### Identification of Differentially Expressed Genes

A generalized linear model identified a total of 12,296 *D. pulex* protein-coding genes across samples with 9% (1,213 genes) identified as differentially expressed (DE) based on chosen threshold criteria (p-value < 0.05, log_2_ fold change of 2) (Table S2).

Collectively, DE genes specific to main effects accounted for ∼30% (total 368 DE genes) of the response while DE genes associated with interaction terms accounted for ∼70% (total 845 DE genes). Among the main effects, phosphorus limitation impacted the greatest number of genes (295), followed by arsenic (46) and then food restriction (27) (Fig. 1A). Few, ∼35%, of main effects DE genes (294 DE genes) were shared among interaction DE gene sets (845 DE genes)(Fig. 1). Annotations for main effects genes can be found in the supplemental materials (Tables S3-5). Interactions gene sets are shown in Figure 1B-D and annotations for DE genes associated with interactions are provided in the supplemental materials (Tables S6-8).

### GO Functional Enrichment of Main Effects and Interactions

The gene ontologies of the DE genes revealed distinct functional differences between main effects and interaction DE gene sets complimenting the magnitude shift in gene expression in the presence of co-stressors. Main effects enrichments revealed specific functional signatures: arsenic broadly impacted structural constituents, enriching terms such as extracellular matrix structural constituent and structural molecule activity (Fig. 2A); low food specifically targeted nutrient uptake and lipid transport, enriching nutrient reservoir activity, lipid transport, and fatty acid transport (Fig. 2C); while low phosphorus enriched diverse regulatory functions and processes— including DNA-binding transcription factor activity, DNA-templated transcription, and Skp-Cullin-F-box containing complex (SCF complex) components—and notably exhibited up to 5X more genes enriched per ontology than the other main effects (Fig. 2B). In contrast, enrichment analysis of interaction DE gene sets revealed an increase in enriched terms by gene count and specificity to the interaction. The low phosphorus × arsenic interaction uniquely enriched chitin biosynthesis and RNA/DNA-mediated processes, alongside developmental processes and regulation of transcription by RNA polymerase II (Fig. 3A), while the low food × low phosphorus interaction enriched various binding and catalytic activities localized to the extracellular region and ER lumen (Fig. 3B). Enrichment could not be assessed for the low food × arsenic interaction due to insufficient DE genes.

**Figure 2:**
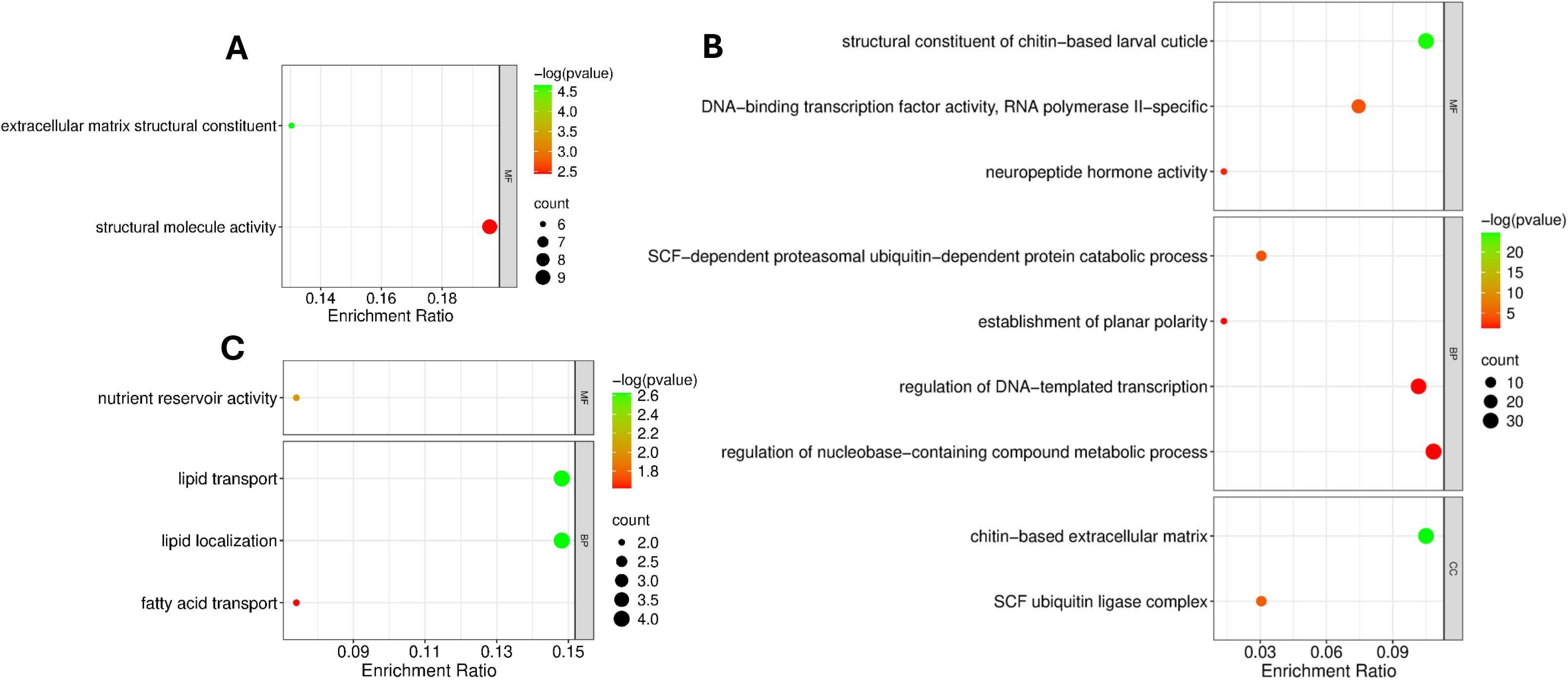
Gene Ontology (GO) enrichment indicates that each main effect enriches distinct GO terms. GO functional enrichment analysis was conducted using gProfiler (g:GOSt) across molecular function (MF, top), biological process (BP, middle), and cellular component (CC, bottom) ontologies for arsenic (A), phosphorus (B), and food (C). A one-tailed Fisher’s test was used to derive P-values with terms being enriched if < 0.05 and corrected for multiple testing (Benjamini &Hochberg, Y; 1995). Enrichment ratio is calculated from the following gProfiler parameters: intersection size/query size.

**Figure 3:**
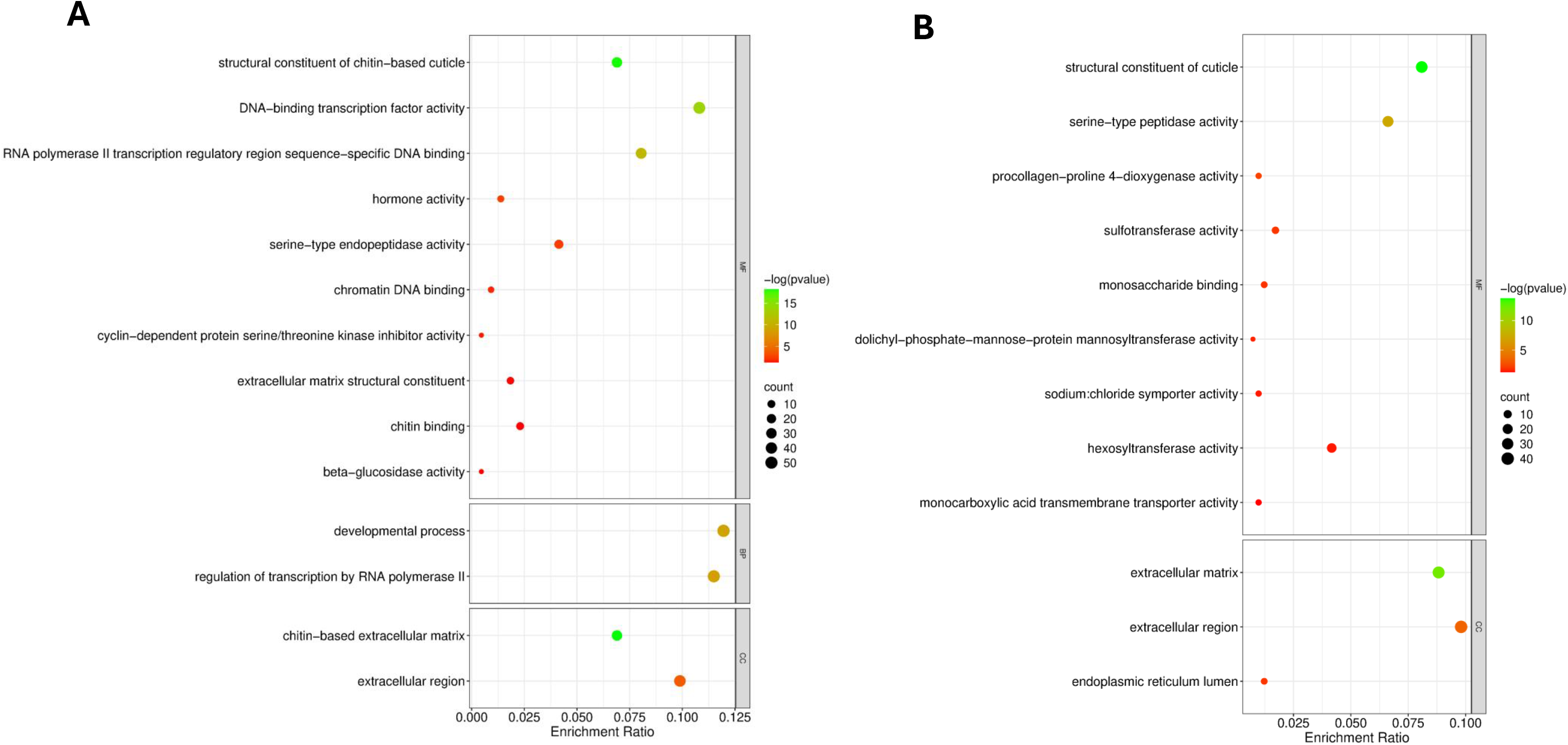
Gene Ontology (GO) enrichment indicate that enriched terms are conditional on low phosphorus x co-stressor. GO functional enrichment analysis was conducted using gProfiler (g:GOSt) across molecular function (MF, top), biological process (BP, middle), and cellular component (CC, bottom) ontologies for arsenic (A), phosphorus x arsenic and (B) low food x low phosphorus. A one-tailed Fisher’s test was used to derive P-values with terms being enriched if < 0.05 and corrected for multiple testing (Benjamini &Hochberg, Y; 1995). Enrichment ratio is calculated from the following gProfiler parameters: intersection size/query size.

### PAA Results on Main Effects and Arsenic x Nutrient Interactions

Pathway activation analysis (PAA) was used to assess directional regulation of KEGG biomolecular pathways by considering the expression levels within enriched pathways of all genes with a measurable log fold change for the effect of interest.

Each gene was classified as induced (positive fold change) or repressed (negative fold change), and a binomial test determined whether the proportion of induced versus repressed genes within a pathway deviated significantly from the null expectation of 0.5 (Hampton et al., 2018). A pathway was reported as significantly activated or repressed only if it contained more than four genes that met the criteria of false discovery rate (FDR) < 0.05 and median log fold change > 1.

PAA revealed substantial differences in the number of activated and repressed pathways across main effects and co-stressor interactions (Table 1; Table S9). Arsenic induced a total of 7 total pathways (4 activated, 3 repressed) activating pathways involved in cell fate and proliferation, while repressing pathways like ascorbate and aldarate metabolism (Tables S9). Phosphorus limitation induced a total of 19 pathways with the majority being activated (14) while the remaining (5) were repressed. Activated pathways include calcium signaling, Hippo signaling, while repressed pathways include metabolism of xenobiotics by cytochrome P450 and steroid hormone biosynthesis (Table S9). Low food induced 10 pathways (5 activation, 5 repressed) including activation of protein digestion and absorption pathways alongside repression of cell cycle, oocyte meiosis, and progesterone-mediated oocyte maturation (Table S9). Low food x low phosphorus repressed a total of 7 pathways including nucleotide excision repair, DNA replication, and cell cycle (Table 1). Low phosphorus x arsenic induced a total of 43 pathways (22 activated, 21 repressed). This interaction activated pathways related to detoxification such as glutathione metabolism and drug metabolism by cytochrome P50. Repressed pathways included base excision repair, spliceosome, mRNA surveillance, and ATP-dependent chromatin remodeling, indicating genotoxic impacts (Table 1). Low food x arsenic induced a total of 5 pathways with 80% being repressed. Repressed pathways including Hippo signaling and various types of N-glycan biosynthesis (Table 1).

### Visualizing Gene Expression Patterns to Identify Synergistic or Antagonistic Interactions

Assessing gene expression patterns can identify whether interacting stressor effects are antagonistic or synergistic (Garcia-Reyero et al., 2012; Shaw et al., 2014; Altschuler et al., 2015, Hampton et al., 2018). Similar to previous studies (Shaw et al., 2014), we visualized expression levels of each DE gene set for a specific treatment, e.g., arsenic, in all treatments including the comprised binary interaction, e.g. arsenic, low phosphorus, and low phosphorus x arsenic. This was completed for all main effects and interactions though for the present study we highlight interactions involving arsenic (Fig. 4; Fig. S4). The magnitude of DE genes for low food (up-regulated genes = 15, down-regulated genes = 12) and arsenic (up-regulated = 13, down-regulated = 33) versus low phosphorus (up-regulated genes = 95, down-regulated genes = 200) is distinct, however, the direction of the fold change remains relatively the same when comparing across main effects suggesting that the expression of these genes remained consistent between main effects (Fig. 4 A, B, D, E; Fig. S4 A, B). In sharp contrast, all arsenic x nutrient interactions exhibited universal antagonism in gene expression responses (Fig. 4, Fig. S4A-C). Both interactive effects of low food x arsenic (down-regulated genes = 2) and low phosphorus x arsenic (up-regulated genes = 225, down-regulated genes = 210) produced a less than additive log fold change; thus, producing an antagonistic response across all differentially expressed genes (Fig. 4C&F).

**Figure 4:**
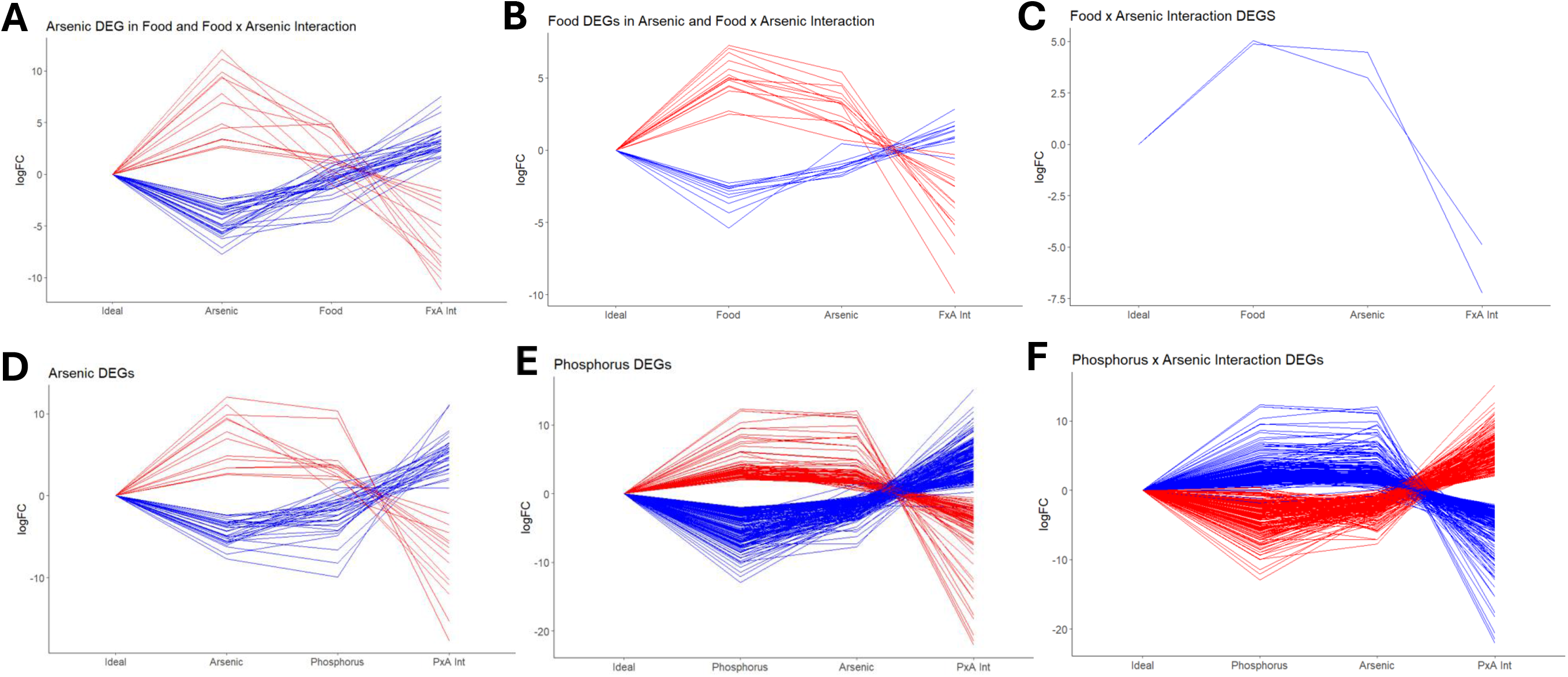
Both arsenic x nutrient interactions, low food x arsenic (A-C) and low phosphorus x arsenic (D-F), produce an antagonistic gene expression response. Differentially expressed (DE) genes for the three-factor linear model that includes the presence of arsenic, low phosphorus quality and low food quantity and their interactions. DE genes were defined by FDR < 0.05 and log_2_fold change > 2 for at least one treatment. DE genes from that treatment are labelled up-regulated (red) or down-regulated (blue). The y-axis represents the log_2_fold change of the DE gene set of interest across the remaining effects. For main effects including arsenic DE genes (A & D), low food DE genes (B), and low phosphorus (D), the plots will exhibit the regulation of DE gene set of interest from left to right; whereas, interaction terms including low food x arsenic (C) and low phosphorus x arsenic (F) will exhibit DE genes regulation from right to left. All subplots of gene expression show a similar pattern of gene regulation being maintained between main effects and the interactions deviate from the expectation of additivity and behave antagonistically.

## Discussion

Natural and anthropogenic environmental stressors most often occur in combination and induce adverse responses in organisms (Folt et al. 1999, Crain et al. 2008, Garcia Reyero et al. 2012, Piggott et al. 2015, Côté et al. 2016). While the standard approach to dissecting interactive effects is to conduct life history assessments and measure impacts on behavior, growth, life span, and reproduction (Folt et al. 1999, Lind et al. 2017, Awoyemi et al. 2020, Schultz et al. 2024); they cannot provide information on how co-stressors impact molecular functions (Roy Chowdhury et al., 2014). Assessing gene expression changes can be utilized to understand an organism’s initial response to the interacting stressors (Shaw et al. 2007; Poynton et al. 2008; Maes et al. 2013). In addition, genome expression studies have been used to understand co-stressor responses in organisms experiencing diverse co-stressor interactions (Shaw et al., 2014; Altschuler et al., 2015; Hampton et al., 2018; Asselman et al., 2019).

In the present study, we assessed the impact of arsenic x nutrient (low food quantity and low phosphorus quality) interactions on gene expression in *D*. *pulex*. Differential gene expression analysis provided gene sets for main-effects and interactions of stressors, while investigations of functional enrichment and pathway activation helped characterize molecular functions. Gene expression patterns were further evaluated to determine whether co-stressor interactions were synergistic or antagonistic. Enrichment analysis revealed the complex interplay of interactions with low phosphorus suggesting that this limiting nutrient is a dominant stressor at the transcriptional level. Pathway activation analysis revealed that each interaction exhibits unique patterns of activating/repressing biomolecular pathways further suggesting that the presence of two stressors creates a conflict in resource allocation and molecular signaling (Bochdanovits and de Jon 2003; Hampton et al., 2018). Assessment of differential gene expression across main effects that compromise an interaction determined that all arsenic x nutrient interactions are antagonistic in nature. Results from the present study support a growing body of evidence that arsenic behaves antagonistically as a co-stressor at varying concentrations, in conjunction with various stressor types, and across multiple different organisms (Shaw et al., 2014; Hampton et al. 2018; Asselman et al., 2019; Ali et al. 2020).

### Differential Expression Reveals Phosphorus Dominance and Substantial Increase in DE for Arsenic x Nutrient Interactions

The differential gene expression patterns observed across main effects revealed distinct, transcriptional signatures: a subtle arsenic effect, a dominant low phosphorus response, and a minimal low food response (Fig. 1). Overlap among main effect DE genes was minimal, suggesting each stressor effect elicits a unique response (Fig.1A). The low arsenic response (46 DE genes) confirms the experimental design, which sought to focus on an environmentally relevant, sublethal arsenic concentration (Schultz et al., 2024), as arsenic alone does not overwhelm the transcriptional landscape. This low level of DE genes aligns with previous work that also observed few DE genes following exposure to low concentrations of arsenic, e.g., in *Daphnia magna* exposed to environmentally relevant arsenic concentration (50 µg L^−1^) which reported a total of 12 arsenic DE genes (FDR < 0.05) (Asselman et al., 2019), and in killifish exposed to arsenic (100 µg L^−1^) where no differential expression was detected (Hampton et al., 2018). Low phosphorus produced a substantially larger DE response (295 genes), suggesting this main effect creates a greater functional consequence than arsenic or low food (Fig. 1A). Similar findings in *D. magna* reported that phosphorus limitation provoked a more widespread transcriptomic response than nitrogen limitation, particularly in metabolic pathways regulating physiological changes (Xu et al., 2021). Low food produced the fewest DE genes (27) (Fig. 1B); however, under identical experimental conditions, starvation had a profound impact on life span, adult growth rate, size and age at maturity in *D. pulex* (Schultz et al., 2024).

Assessment of interactions revealed distinct transcriptional responses, with the most dramatic contrast emerging between the arsenic x nutrient interactions (Fig. 1B&D). Co-stressor interactions involving low phosphorus (low food x low phosphorus and low phosphorus x arsenic) experienced a substantial increase in differential expression in comparison to main effect gene sets. The pattern of increased differential expression is commonly observed in co-stressor interactions (Yang et al., 2007; Vandenbrouck et al., 2009), however; the magnitude difference in arsenic x nutrient interactions differed greatly. The low phosphorus × arsenic interaction produced 435 DE genes (nearly 10x the arsenic and 1.5x the phosphorus main effects) whereas the low food × arsenic interaction yielded only 2 DE genes. Genes uniquely induced under low phosphorus × arsenic included several cuticle proteins and digestive enzymes (Table S6), suggesting specific physiological impacts on molting, structural integrity, and nutrient assimilation which have been reported under low phosphorus (Jeyasingh et al., 2011; Roy Chowdhury et al., 2014) and arsenic condition, respectively (Bi et al., 2024; Lin et al, 2024). These findings provide important implications for understanding how nutrient availability shapes organismal sensitivity to environmental contaminants.

### GO Results Suggest Diverse Biological Effects from Arsenic x Nutrient Interactions in Comparison to Main Effects

Enrichment analyses indicate that each main effect impacts distinct KEGG biochemical pathways (Fig. 2) while interactive effects exhibit a complex interplay of stress responses (Fig. 3). Enriched genes for arsenic were broad and only impacted the extracellular matrix structural constituents and structural molecule activity. The response observed also remains consistent with the design, which featured a sublethal, environmentally relevant concentration of arsenic (Fig. 3A; Schultz et al., 2024). Low food, by contrast, enriched specific terms related to lipid storage and transport, including nutrient reservoir activity, lipid transport, and fatty acid transport (Fig. 3C). It has been previously reported that adult *Daphnia* are more reliant on lipids for egg production (Goulden & Henry 1984, Muller Navarra 1995). These enriched genes parallel observations from the companion life history study and could potentially explain why low food significantly impacted adult *D. pulex* growth rate in comparison to phosphorus limitation (Schultz et al., 2024). Low phosphorus enriched more unique terms across ontologies and at a larger scale (30 genes) in comparison to arsenic and low food (2-9 genes)(Fig. 2B). Terms such as structural constituent of chitin-based larval cuticle functions and DNA-mediated processes such as DNA-binding transcription factor activity and DNA-templated transcription were enriched under phosphorus limitation.

These results are plausible considering chitin biosynthesis and modification, as well as DNA-mediated pathways are inherently phosphorus-dependent processes (Yang and Fukamizo 2019) and are critical for appropriate development in invertebrates (Bi et al., 2024; Lin 2024). Additionally, the enrichment of both the SCF complex and SCF-dependent catabolic process reflects a cellular quality control response to prepare for proteasomal degradation under phosphorus limitation (Fig. 2B; Thompson et al., 2021).

Enrichment analyses revealed the interactive effects increased both in gene count and functional specificity compared to main effects (Fig. 3). In other words, there was a fundamental difference in how biomolecular pathways were influenced by main effects and interaction DE gene sets. The low phosphorus × arsenic interaction enriched diverse functional terms across binding, catalytic, and structural activities (Fig. 3B). This interaction reported an increase in chitin-related terms as well as DNA-associated regulatory pathways. Previous studies have indicated that phosphorus limitation and arsenic can independently produce similar responses. The expression of cuticular proteins in *D. magna* were altered by arsenic in *D. magna* (Asselman et al. 2019) and *D. pulex* (Roy Chowdhury et al. 2015); strongly suggesting that the interaction of co-stressors would impact the same functions. Additionally, the enrichment of DNA-binding transcription factor activity, chromatin binding, and developmental processes suggests that arsenic coordinately disrupts chromatin organization and DNA-associated regulatory pathways. Arsenic exposure is well known to impair DNA repair and replication, alter chromatin structure, and suppress the expression of genes involved in these processes (Hughes, 2002; Yang & Frenkel, 2002; Tam Price & Wang, 2020; Medda et al., 2021). Increasing transcriptomic evidence indicates that these molecular responses to low levels of arsenic are also conserved in invertebrates (Bi et al., 2024; Lin et al., 2024). Serine-type endopeptidase and hormone activity were also significantly enriched by the low phosphorus x arsenic interaction suggesting impacts on protein digestion as peptidase activity is critical for the growth and development of invertebrate species through the digestion of dietary proteins to be used for protein synthesis (Park and Kwak, 2020). The interaction of low food x low phosphorus suggests potential adverse impacts on essential protein modification processes related to structural cuticle integrity (Fig. 4B). The molecular functions, structural constituent of the cuticle and dolichyl-phosphate-mannose-protein mannosyltransferase activity suggest alterations to protein glycosylation (Girrbach et al., 2000). More specifically, O-glycosylation is an evolutionarily conserved process that is initiated in the endoplasmic reticulum and critical for the stability and localization of proteins (Martin-Blanco & Garcia-Bellido, 1996; Girrbach et al., 2000). Previous studies have reported that disruptions of mannosyltransferase activity reduce viability in organisms such as *S. cerevisiae* (Girrbach et al., 2000) and cause defects in muscle structures and cuticle alignment in *D. melanogaster* (Martin-Blanco & Garcia-Bellido, 1996) implying detrimental impacts on growth and development. In contrast, the low food × arsenic interaction could not be assessed due to insufficient DE genes, a limitation that itself underscores the near absence of a transcriptional response when arsenic is combined with food limitation.

Enrichment analysis reveals that dominance exhibited by low phosphorus is limited to an increase in transcription and is not observed in early functional measures. The increase in differential expression observed under low phosphorus as a main effect and as a result of interacting with additional co-stressor (low phosphorus x arsenic & low food x low phosphorus) would suggest that low phosphorus would not only strongly influence the quantity of genes but also dominate the functional response. Our enrichment results suggest that phosphorus limitation increases DE gene count; however, the functional response is conditional based on the co-stressor (Fig. 3A & 3B).

Collectively, the enriched GO terms suggest that interactive effects produce functionally different responses than the main effects.

### PAA Suggests Shift in Pathway Regulation between Co-stressor Interactions and Main Effects

To further understand the observed differences in complexity and severity, we assessed impacts at the pathway level using PAA. Results from the current study demonstrate three key findings: (1) each main effect disrupts or enables unique KEGG pathways, (2) pathway level results suggests a directional shift in regulation between main effects and interactions, and (3) interactions produce nonadditive responses at the pathway level. However, we acknowledge that further elucidation of these specific pathways is warranted.

PAA revealed that each main effect produced a distinct pathway-level signature (Table S9). The activation of cell proliferation pathways under arsenic exposure aligns with well-documented mechanisms of arsenic toxicity (Hughes 2002, Yang and Frankel 2002, Fan et al., 2015, Ali et al., 2020, Tam Price and Wang 2020) and has been detected previously in killifish at similar arsenic concentrations (Hampton et al. 2018). A previous study reported that alterations in ascorbate and aldarate metabolism under arsenic exposure are indicative of oxidative damage in freshwater snails (Bi et al. 2024). Low phosphorus activated nearly 75% of pathways including critical cell signaling such as calcium and Hippo signaling (Table S9). Taken in conjunction with the significant co-enrichment of the SCF complex specific terms, these results suggest that phosphorus limitation may impact intracellular protein degradation which is an ATP dependent process responsible for regulating signal transduction, cell cycle progression, and gene transcription in eukaryotes (Thompson et al., 2021). This finding is particularly interesting given that arsenic has been experimentally linked to increased ubiquitination through E3 ligase activity in killifish (Shaw et al., 2010). These pathways are critical for growth and development and could plausibly explain the reduction in juvenile growth rate and size at first reproduction observed in Schultz et al., 2024. Low food exhibited an even split of activation and repression across KEGG pathways associated with nutrient acquisition and reproduction (Table S9). Promotion of nutrient acquisition while suppressing reproductive pathways suggests that low food physiologically demands reprioritization of resource allocation toward survival rather than reproduction. Life-history tradeoffs can manifest at the transcriptomic level as a shift between energy metabolism and protein biosynthesis, with signaling pathways coordinating this allocation (Bochdanovits & de Jong, 2003).

PAA revealed co-stressor interactions exhibited opposing directionality in comparison to main effects, suggesting a fundamental reprogramming of pathway regulation under co-stressor interactions. Low phosphorus x arsenic interaction shares a collective 15 significant pathways with its constituent main effects and every pathway exhibited opposite directionality under the interaction (Fig. S5). For example, the highly conserved cell signaling pathway, Hippo signaling, was significantly activated within the phosphorus limitation and arsenic DE gene sets (Table S9); however, it was significantly repressed in arsenic x nutrient interaction (Table 1). The shift in biological function is likely due to arsenic and phosphorus sharing similar chemical and biological properties where the co-occurrence results in detrimentally impacting the same pathways (Villa-Bellosta & Sorribas, 2010). For example, both arsenic (Tam Price & Wang, 2020) and phosphorus limitation (Xu et al., 2021) promote cell cycle disruption, the extensive pathway-level reprogramming observed in the low phosphorus x arsenic interaction, corresponds well with documented fitness impacts in *Daphnia* (Awoyemi et al., 2020; Schultz et al., 2024). Low food x arsenic interaction exhibited repression with only one activated pathway (e.g. lysosome pathway) suggesting that energy demands may shift away from growth and reproduction toward survival through enhanced protein degradation (Ellgaard et al., 2003) and aligns with the observation that only low food and arsenic significantly impacted *D. pulex* lifespan (Schultz et al., 2024).

PAA also revealed that interactions with low food impacted fewer pathways in comparison to main effects. It is commonly reported that the combination of two stressors will increase transcriptional response likely leading to more pathways being impacted (Vandenbrouck et al., 2009; Altshuler et al., 2015); however, interactions with low food (low food x arsenic and low food x low phosphorus) exhibited a reduction in response by impacting only 5 and 7 pathways, respectively. Considering the low number of pathways impacted and the majority being repressed, this adds support to the hypothesis suggested by others that starvation prevents the organism from partaking in energy intensive processes (Knops et al., 2001; Villarroel et al., 2009). In contrast, the low phosphorus x arsenic interaction induced a total of 43 pathways which is a significant increase in comparison to the other interactions (Table 1). In comparison to both main effects and other interactions, this significant increase in the total number of pathways impacted is likely due to the chemical similarities between phosphorus and arsenic and competing for uptake in phosphorus dominated biochemical pathways (Villa-Bellosta & Sorribas, 2010; Byeon et al., 2024). Similarly, PAA further reinforced that interactions with low phosphorus exhibit a functional response conditional to the co-stressor and indicating that dominance in differential expression did not translate to functional measures. Determining whether a stressor of interest exhibits dominances over other stressors is critical in the identification and mitigation of co-stressor interactions (Crain et al., 2008 and Côté et al., 2016). A previous study conducted by Folt et al. (1999) reported that the dominant stressor unique to the two species tested (e.g. *D. pulex* and *D. pulicaria*) appeared to drive the magnitude of the stress effect. Our results support the dominant stressor influences the magnitude of response from comparing DE gene sets to pathway level results. The low phosphorus x arsenic interaction produced a broad response in comparison to both quantity (total 43) and type while the low phosphorus x food interaction produced a functionally narrow response in both quantity (total 7) and type of pathways induced (types listed in Table 1). Assessing differential expression as well as more functional measures such as pathway-level impacts provides a more nuanced understanding of the conditional behavior of co-stressor.

### Arsenic x Nutrient Interactions in Gene Expression are Antagonistic in Nature

Pathway activation analysis (PAA) provides critical insight into the systemic impact of arsenic x nutrient interactions by revealing both the magnitude and direction of pathway-level regulation under co-stressor conditions. However, interpreting these activation and repression patterns requires careful consideration of how individual stressors and their interactions deviate from additive expectations. Therefore, evaluating whether these interactions are synergistic or antagonistic is essential to understanding the nature of arsenic x nutrient interactions. A previous meta-analysis suggests that interactions between toxins and nutrients are typically antagonistic in nature (Crain et al., 2008). More specifically, interactions between arsenic and other co-stressors have consistently yielded antagonistic impacts on gene expression in both vertebrate and invertebrate species. For example, a study assessing the influence of arsenic exposure on acclimation to changing salinity across different killifish populations reported uniformly antagonistic responses in gene expression (Shaw et al., 2014; Hampton et al., 2018). Similarly, a genome-wide assessment of natural populations of *Daphnia magna* exposed to arsenic and copper revealed an antagonistic gene expression pattern in critical cuticle proteins (Asselman et al., 2019). In the present study, the visualization of gene expression log_2_fold changes between main effects and interaction DE suggests that all arsenic x nutrient interactions are antagonistic (Fig. 4; Fig. S4). While it is a common prediction that co-stressor combinations do not meet the assumption of additivity and produce nonadditive effects on expression patterns (Yang et al., 2007; Vandenbrouck et al., 2009; Garcia-Revero et al., 2012; Maes et al., 2013; Altshuler et al., 2015), the present study adds to growing body of literature that has observed arsenic x co-stressor interactions at the level of gene expression to be entirely antagonistic (Shaw et al., 2014; Hampton et al., 2018; Asselman et al., 2019).

## Conclusion

This study demonstrates that nutritional status differentially influences the transcriptomic response to arsenic exposure in *D*. *pulex*. The environmentally relevant, arsenic concentration used in this study’s design was strategically chosen to interrogate how nutrient co-stressors modulate arsenic response. A consistent pattern emerged across gene and pathway levels: phosphorus limitation exerted a dominant influence both as a main effect and as a co-stressor, transcriptional responses to interactions were largely unique to each stressor combination, and low food quantity and low phosphorus concentrations modulated arsenic toxicity through distinct molecular mechanisms. Notably, the contrast between the robust low phosphorus × arsenic interaction and the minimal low food × arsenic interaction underscores the specificity of phosphorus limitation in mediating arsenic toxicity, likely through competitive uptake and shared metabolic pathways. Our results reveal that arsenic x nutrient interactions are uniformly antagonistic at the level of gene expression, supporting the growing body of literature suggesting that arsenic behaves uniquely, antagonistically as a co-stressor.

This antagonism suggests that nutrient limitation does not simply amplify arsenic toxicity but instead redirects transcriptional priorities, likely compromising fitness through mechanisms distinct from arsenic exposure alone. This study highlights a critical gap in standard ecological risk assessment (ERA), which typically evaluates contaminants in isolation without accounting for nutritional status. Our findings demonstrate that co-stressor interactions can fundamentally alter transcriptional responses in ways that cannot be predicted from single-stressor data alone. Incorporating transcriptomic assessment of nutrient-contaminant interactions into ERA frameworks would improve predictive capacity for real-world exposures, where organisms face complex mixtures of stressors. The companion fitness study using this same experimental design (Schultz et al., 2024) strengthens our transcriptional findings, yet mechanistic gaps remain in linking these molecular events to the reported life history changes. Future efforts should prioritize experimentally connecting these key events to strengthen AOP development and support the adoption of new approach methodologies (NAMs). Given the prevalence of arsenic in the environment, understanding arsenic x nutrient interactions is essential for protecting aquatic ecosystems and the human populations that depend on them.

## Data Availability

The raw Next-seq RNA sequence and processed data from this study have been uploaded to the NCBI Gene Expression Omnibus Accession GSE341878.

## Acknowledgements

We are grateful to B. Jackson and the Dartmouth Trace Element Core Facility for arsenic measurements. We also thank Kelley Thomas, Joseph Sevygny and the Hubard Center for Genome Studies at the University of New Hampshire for library preparation and RNA sequencing. Additionally, we thank Emilyann Ashford and Autumn Berlied for their help in maintaining the experimental cultures.

## Author Contributions

Emily R. DeTemple: Data curation; Data Analysis; Writing – original draft; Writing – review & editing. Craig E. Jackson: Data curation; Writing – reviewing & editing. Anthony Schultz: Maintenance of experimental cultures; RNA isolation. Thomas H. Hampton: Supervision; Writing – Review & Editing. Joseph R. Shaw: Supervision; Writing – Review & Editing. Priyanka Roy Chowdhury: Conceptualization; Funding acquisition; Methodology; Project administration; Resources; Supervision; Writing-Review & Editing.

## References

1. Agency for Toxic Substances and Disease Registry. (2026, April). Support document to the 2025 substance priority list (candidates for toxicological profiles). U.S. Department of Health and Human Services. https://www.atsdr.cdc.gov/spl/resources/ATSDR-2025-SPL-Support-Document-508.pdf

2. Allen, H. J., Impellitteri, C. A., Macke, D. A., Heckman, J. L., Poynton, H. C., Lazorchak, J. M., … & Nadagouda, M. N. (2010). Effects from filtration, capping agents, and presence/absence of food on the toxicity of silver nanoparticles to Daphnia magna. Environmental Toxicology and Chemistry, 29(12), 2742–2750.

3. Ali, W., Zhang, H., Junaid, M., Mao, K., Xu, N., Chang, C., Rasool, A., Wajahat Aslam, M., Ali, J., & Yang, Z. (2021). Insights into the mechanisms of arsenic-selenium interactions and the associated toxicity in plants, animals, and humans: A critical review. Critical Reviews in Environmental Science and Technology, 51(7), 704–750. 10.1080/10643389.2020.1740042

4. Altshuler, I., McLeod, A. M., Colbourne, J. K., Yan, N. D., & Cristescu, M. E. (2015). Synergistic interactions of biotic and abiotic environmental stressors on gene expression. Genome, 58(3), 99–109. 10.1139/gen-2015-0045

5. Amtmann, A., & Armengaud, P. (2009). Effects of N, P, K and S on metabolism: new knowledge gained from multi-level analysis. Current opinion in plant biology, 12(3), 275–283.

6. Ashburner, M., Ball, C. A., Blake, J. A., Botstein, D., Butler, H., Cherry, J. M., Davis, A. P., Dolinski, K., Dwight, S. S., Eppig, J. T., Harris, M. A., Hill, D. P., Issel-Tarver, L., Kasarskis, A., Lewis, S., Matese, J. C., Richardson, J. E., Ringwald, M., Rubin, G. M., & Sherlock, G. (2000). Gene Ontology: Tool for the unification of biology. Nature Genetics, 25(1), 25–29. 10.1038/75556

7. Asselman, J., Semmouri, I., Jackson, C., Keith, N., Van Nieuwerburgh, F., Deforce, D., Shaw, J., & Schamphelaere, K. (2019). Genome-Wide Stress Responses to Copper and Arsenic in a Field Population of Daphnia. Environmental Science and Technology, 53, 3850–3859.

8. Asselman, J., De Coninck, D. I., Beert, E., Janssen, C. R., Orsini, L., Pfrender, M. E., Decaestecker, E., & De Schamphelaere, K. A. (2017). Bisulfite Sequencing with *Daphnia* Highlights a Role for Epigenetics in Regulating Stress Response to *Microcystis* through Preferential Differential Methylation of Serine and Threonine Amino Acids. Environmental Science & Technology, 51(2), 924–931. 10.1021/acs.est.6b03870

9. Awoyemi, O., Subbiah, S., Thompson, K., Velazquez, A., Peace, A., & Mayer, G. (2020). Trophic-Level Interactive Effects of Phosphorus Availability on the Toxicities of Cadmium, Arsenic, and their Binary Mixture in Media-Exposed Scenedesmus acutus and Media and Dietary-Exposed Daphnia pulex. Environmental Science and Technology, 54, 5651–5666.

10. Ayotte, J. D., Montgomery, D. L., Flanagan, S. M., & Robinson, K. W. (2003). Arsenic in groundwater in eastern New England: occurrence, controls, and human health implications. Environmental science & technology, 37(10), 2075–2083.

11. Becker, D., Reydelet, Y., Lopez, J. A., Jackson, C., Colbourne, J. K., Hawat, S., … & Paul, R. J. (2018). The transcriptomic and proteomic responses of Daphnia pulex to changes in temperature and food supply comprise environment-specific and clone-specific elements. BMC genomics, 19(1), 376.

12. Benjamini, Y., & Hochberg, Y. (1995). Controlling the false discovery rate: A practical and powerful approach to multiple testing. J R Stat Soc Ser B Stat Methodol, 57(1), 289–300.

13. Bi, X., Qiu, M., Li, D., Zhang, Y., Zhan, W., Wang, Z., Lv, Z., Li, H., & Chen, G. (2024). Transcriptomic and metabolomic analysis of the mechanisms underlying stress responses of the freshwater snail, *Pomacea canaliculata*, exposed to different levels of arsenic. Aquatic Toxicology, 267, 106835. 10.1016/j.aquatox.2024.106835

14. Bochdanovits, Z., & de Jong, G. (2004). Antagonistic pleiotropy for life history traits at the gene expression level. The Royal Society, 271, S75–S78.

15. Boersma, M., & Kreutzer, C. (2002). Life at the edge: Is food quality really of minor importance at low quantities? Ecology, 83(9), 2552–2561.

16. Bolger, A. M., Lohse, M., & Usadel, B. (2014). Trimmomatic: A flexible trimmer for Illumina sequence data. Bioinformatics, 30(15), 2114–2120. 10.1093/bioinformatics/btu170

17. Borgono, J. M., Vicent, P., Venturino, H., & Infante, A. (1977). Arsenic in the drinking water of the city of Antofagasta: Epidemiological and clinical study before and after the installation of the treatment plant. Environ mental Health

18. Breitburg, D.L., Baxter, J.W., Hatfield, C.A. (1998). Understanding effects of multiple stressors: ideas and challenges. In: Successes, Limitations, and Frontiers in Ecosystem Science (eds Pace, M.L. & Groffman, P.M.). Springer, New York, pp. 416–431.

19. Byeon, E., Kang, H.-M., Yoon, C., & Lee, J.-S. (2021). Toxicity mechanisms of arsenic compounds in aquatic organisms. Aquatic Toxicology, 237, 105901. 10.1016/j.aquatox.2021.105901

20. Cheng, Y., Ding, J., Arenas, C. E. D., Brinkmann, M., & Ji, X. (2024). A brief review on the assessment of potential joint effects of complex mixtures of contaminants in the environment. Environmental Science: Advances, 3(5), 661–675.

21. Côté, I. M., Darling, E. S., & Brown, C. J. (2016). Interactions among ecosystem stressors and their importance in conservation. Proceedings of the Royal Society B: Biological Sciences, 283(1824), 20152592. 10.1098/rspb.2015.2592

22. Crain, C. M., Kroeker, K., & Halpern, B. S. (2008). Interactive and cumulative effects of multiple human stressors in marine systems. Ecology Letters, 11(12), 1304–1315. 10.1111/j.1461-0248.2008.01253.x

23. Dodds, W. K. (2002). Freshwater ecology: Concepts and environmental applications. Academic Press.

24. Dreval, K., Tryndyak, V., Kindrat, I., Twaddle, N.C., Orisakwe, O.E., Mudalige, T.K., Beland, F.A., Doerge, D.R. and Pogribny, I.P., 2018. Cellular and molecular effects of prolonged low-level sodium arsenite exposure on human hepatic HepaRG cells. Toxicological Sciences, 162(2), pp.676–687.

25. Ellgaard, L., & Helenius, A. (2003). Quality control in the endoplasmic reticulum. Nature Reviews Molecular Cell Biology, 4(3), 181–191. 10.1038/nrm1052

26. Embree, C. M., Paul, D., Stephanou, A., & Singh, G. (2025). Direct and indirect effects of spliceosome disruption compromise gene regulation by nonsense-mediated mRNA decay. RNA Biology, 22(1), 1–26. 10.1080/15476286.2025.2552517

27. Eustermann, S., Patel, A. B., Hopfner, K.-P., He, Y., & Korber, P. (2024). Energy-driven genome regulation by ATP-dependent chromatin remodellers. Nature Reviews Molecular Cell Biology, 25(4), 309–332. 10.1038/s41580-023-00683-y

28. Fan, W., Ren, J., Li, X., Wei, C., Xue, F., & Zhang, N. (2015). Bioaccumulation and oxidative stress in *Daphnia magna* exposed to arsenite and arsenate. Environmental Toxicology and Chemistry, 34(11), 2629–2635. 10.1002/etc.3119

29. Folt, C. L., Chen, C. Y., Moore, M. V., & Burnaford, J. (1999). Synergism and antagonism among multiple stressors. American Society of Limnology and Oceanography, 44, 864–877.

30. Garcia-Reyero, N., Escalon, B. L., Loh, P., Laird, J. G., Kennedy, A. J., Berger, B., & Perkins, E. J. (2012). Assessment of Chemical Mixtures and Groundwater Effects on *Daphnia magna* Transcriptomics. Environmental Science & Technology, 46(1), 42–50. 10.1021/es201245b

31. Girrbach, V., Zeller, T., Priesmeier, M., & Strahl-Bolsinger, S. (2000). Structure-Function Analysis of the Dolichyl Phosphate-Mannose: Protein O-Mannosyltransferase ScPmt1p*. 275, 19288–19296.

32. Goulden, C. E., & Henry, L. L. (1984). 7. Lipid energy reserves and trophic interactions within aquatic ecosystems, 85, 167.

33. Hampton, T., Jackson, C., Jung, D., Chen, C., Glaholt, S., Stanton, B., Colbourne, J., & Shaw, J. (2018). Arsenic Reduces Gene Expression Response to Changing Salinity in Killifish. Environmental Science and Toxicology, 52, 8811–8821.

34. Hennig, B., Ormsbee, L., McClain, C. J., Watkins, B. A., Blumberg, B., Bachas, L. G., Sanderson, W., Thompson, C., & Suk, W. A. (2012). Nutrition Can Modulate the Toxicity of Environmental Pollutants: Implications in Risk Assessment and Human Health. Environmental Health Perspectives, 120(6), 771–774. 10.1289/ehp.1104712

35. Hilleren, P., & Parker, R. (1999). Mechanisms of mRNA surveillance in eukaryotes. Annual Review of Genetics, 33, 229–260.

36. Hong, K. H., Keen, C. L., Mizuno, Y., Johnston, K. E., & Tamura, T. (2000). Effects of dietary zinc deficiency on homocysteine and folate metabolism in rats. Journal of Nutritional Biochemistry, 11(3), 165–169. 10.1016/s0955-2863(99)00089-3

37. Hsueh, Y. M., Cheng, G. S., Wu, M. M., Kuo, T. L., & Chen, C. J. (1995). Multiple risk factors associated with arsenic-induced skin cancer: Effects of chronic liver diseases and malnutritional status. British Journal of Cancer, 71(1), 109–114. 10.1038/bjc.1995.22

38. Hughes, M. (2002). Arsenic toxicity and potential mechanisms. Toxicology Letters, 133, 1–16.

39. International Agency for Research on Cancer (IARC). (1980). IARC mono graphs on the evaluation of the carcinogenic risk of chemicals to hu mans: Some metals and metallic compounds (Vol. 23).

40. Jeyasingh, P. D., Goos, J. M., Lind, P. R., Roy Chowdhury, P., & Sherman, R. E. (2020). Phosphorus supply shifts the quotas of multiple elements in algae and *Daphnia*: Ionomic basis of stoichiometric constraints. Ecology Letters, 23(7), 1064–1072. 10.1111/ele.13505

41. Jeyasingh, P. D., Ragavendran, A., Paland, S., Lopez, J. A., Sterner, R. W., & Colbourne, J. K. (2011). How do consumers deal with stoichiometric constraints? Lessons from functional genomics using Daphnia pulex: GENOMIC RESPONSE TO STOICHIOMETRIC IMBALANCE. Molecular Ecology, 20(11), 2341–2352. 10.1111/j.1365-294X.2011.05102.x

42. Kaamoush, M., & El-Agawany, N. (2026). Nutrient metal interactions and adaptive responses of Dunaliella tertiolecta to zinc and copper toxicity under phosphorus limitation. Scientific Reports, 16(1), 13399. 10.1038/s41598-026-47929-1

43. Kilham, S. S., Kreeger, D. A., Lynn, S. G., Goulden, C. E., & Herrera, L. (1998). COMBO: A defined freshwater culture medium for algae and zooplankton. Hydrobiologia, 377(1-3), 147–159. 10.1023/A:1003231628456

44. Kim, H.J., Koedrith, P., Seo, Y.R. (2015) Ecotoxicogenomic Approaches for Understanding Molecular Mechanisms of Environmental Chemical Toxicity Using Aquatic Invertebrate, Daphnia Model Organism. Gu J-D, ed. International Journal of Molecular Sciences 16(6):12261–12287.

45. Knops, M., Altenburger, R., & Segner, H. (2001). Alterations of physiological energetics, growth, and reproduction of Daphnia magna under toxicant stress. Aquatic Toxicology, 53(2), 79–90.

46. Kolberg, L., Raudvere, U., Kuzmin, I., Adler, P., Vilo, J., & Peterson, H. (2023). g:Profiler—Interoperable web service for functional enrichment analysis and gene identifier mapping (2023 update). Nucleic Acids Research, 51(W1), W207–W212. 10.1093/nar/gkad347

47. Kooijman, S. A. L. M. (2000). Dynamic energy and mass budgets in bio logical systems. Cambridge University Press.

48. Lampert, W. (1977). Studies on the carbon balance of Daphnia pulex de Geer as related to environmental conditions. 4. Determination of the threshold concentration as a factor controlling the abundance

49. Li, L., Ekström, E. C., Goessler, W., Lönnerdal, B., Nermell, B., Yunus, M., … & Vahter, M. (2008). Nutritional status has marginal influence on the metabolism of inorganic arsenic in pregnant Bangladeshi women. Environmental health perspectives 116(3): 315.

50. Lin, X., Wang, W., & He, F. (2024). Molecular level toxicity effects of As(V) on Folsomia candida: Integrated transcriptomics and metabolomics analyses. Science of the Total Environment, 922.

51. Lind, P. R., & Jeyasingh, P. D. (2018). Interactive effects of dietary phosphorus and iron on *Daphnia* life history. Limnology and Oceanography, 63(3), 1181–1190. 10.1002/lno.10763

52. Maes, G. E., Raeymaekers, J. A. M., Hellemans, B., Geeraerts, C., Parmentier, K., De Temmerman, L., Volckaert, F. A. M., & Belpaire, C. (2013). Gene transcription reflects poor health status of resident European eel chronically exposed to environmental pollutants. Aquatic Toxicology, 126, 242–255. 10.1016/j.aquatox.2012.11.006

53. Martin-Blanco, E., & Garcia-Bellido, A. (1996). Mutations in the rotated abdomen locus affect muscle development and reveal an intrinsic asymmetry in Drosophila. Proc Natl Acad Sci U S A, 93, 6048–6052.

54. Mazumder, D. N. G., Haque, R., Ghosh, N., De, B. K., Santra, A., Chak raborty, D., & Smith, A. H. (1998). Arsenic levels in drinking water and the prevalence of skin lesions in West Bengal, India. International Journal of Epidemiology, 27(5), 871–877. 10.1093/ije/27.5.871

55. Medda, N., De, S. K., & Maiti, S. (2021). Different mechanisms of arsenic related signaling in cellular proliferation, apoptosis and neo-plastic transformation. Ecotoxicology and Environmental Safety, 208, 111752. 10.1016/j.ecoenv.2020.111752

56. Meharg, A. A., & Hartley-Whitaker, J. (2002). Arsenic uptake and metabolism in arsenic resistant and nonresistant plant species. New Phytologist, 154(1), 29–43.

57. Miao, A.-J., Wang, N.-X., Yang, L.-Y., & Wang, W.-X. (2012). Accumulation kinetics of arsenic in *Daphnia magna* under different phosphorus and food density regimes. Environmental Toxicology and Chemistry, 31(6), 1283–1291. 10.1002/etc.1822

58. Park, K., & Kwak, I.-S. (2020). Cadmium-induced developmental alteration and upregulation of serine-type endopeptidase transcripts in wild freshwater populations of Chironomus plumosus. Ecotoxicology and Environmental Safety, 192, 110240. 10.1016/j.ecoenv.2020.110240

59. Partridge, L., Piper, M. D., & Mair, W. (2005). Dietary restriction in Drosophila. Mechanisms of ageing and development, 126(9), 938–950.

60. Patro, R., Duggal, G., Love, M. I., Irizarry, R. A., & Kingsford, C. (2017). Salmon provides fast and bias-aware quantification of transcript expression. Nature Methods, 14(4), 417–419. 10.1038/nmeth.4197

61. Piggott, J. J., Townsend, C. R., & Matthaei, C. D. (2015). Reconceptualizing synergism and antagonism among multiple stressors. Ecology and Evolution, 5(7), 1538–1547. 10.1002/ece3.1465

62. Poynton, H. C., Loguinov, A. V., Varshavsky, J. R., Chan, S., Perkins, E. J., & Vulpe, C. D. (2008). Gene Expression Profiling in *Daphnia magna* Part I: Concentration-Dependent Profiles Provide Support for the No Observed Transcriptional Effect Level. Environmental Science & Technology, 42(16), 6250–6256. 10.1021/es8010783

63. Qiu, T., Tao, Y., Yao, X., Wang, N., Jiang, L., & Sun, X. (2026). Arsenic Methyltransferase Function in Inorganic Arsenic Biotransformation: Implications for Health Risks and Disease Development. Biological Trace Element Research, 1-12.

64. Quirós, L., Piña, B., Solé, M., Blasco, J., López, M. Á., Riva, M. C., Barceló, D., & Raldúa, D. (2007). Environmental monitoring by gene expression biomarkers in Barbus graellsii: Laboratory and field studies. Chemosphere, 67(6), 1144–1154. 10.1016/j.chemosphere.2006.11.032

65. Ravenscroft, P., Brammer, H., & Richards, K. (2009). Arsenic Pollution: A Global Synthesis (1st ed.). Wiley. 10.1002/9781444308785

66. Rhind, S. M. (2009). Anthropogenic pollutants: A threat to ecosystem sustainability? Philosophical Transactions of the Royal Society, B: Biological Sciences, 364(1534), 3391–3401.

67. Robinson, M. D., McCarthy, D. J., & Smyth, G. K. (2010). edgeR: A Bioconductor package for differential expression analysis of digital gene expression data. Bioinformatics, 26(1), 139–140. 10.1093/bioinformatics/btp616

68. Roy Chowdhury, P., Frisch, D., Becker, D., Lopez, J. A., Weider, L. J., Colbourne, J. K., & Jeyasingh, P. D. (2015). Differential transcriptomic responses of ancient and modern *Daphnia* genotypes to phosphorus supply. Molecular Ecology, 24(1), 123–135. 10.1111/mec.13009

69. Roy Chowdhury, P., Lopez, J., Weider, L., Colbourne, J., & Jeyasingh, P. (2014). Functional Genomics of Intraspecific Variation in Carbon and Phosphorus Kinetics in Daphnia. Journal of Experimental Zoology, 321, 387–398.

70. Sales, S. C. M., Rietzler, A. C., & Ribeiro, M. M. (2016). Arsenic toxicity to cladocerans isolated and associated with iron: Implications for aquatic environments. Anais Da Academia Brasileira de Ciências, 88(suppl 1), 539–548. 10.1590/0001-3765201620140670

71. Schultz, A., Owens, J., Demidenko, E., & Roy Chowdhury, P. (2024). Differential Toxicity of Arsenic in Daphnia pulex Under Phosphorus and Food Limitation. Environmental Toxicology and Chemistry, 43(8), 1807–1819.

72. Shaw, J. R., Dempsey, T. D., Chen, C. Y., Hamilton, J. W., & Folt, C. L. (2006). Comparative toxicity of cadmium, zinc and mixtures of cadmium and zinc to daphnids. Environmental Toxicology and Chemistry, 25(1), 182–189.

73. Shaw, J., Glaholt, S., Greenberg, G., Alvarez, R., & Folt, C. (2007). Acute Toxicity of Arsenic to Daphnia pulex: Influence of Organic Functional Groups and Oxidation State. Environmental Toxicology and Chemistry, 26(7), 1532–1537.

74. Shaw, J. R., Bomberger, J. M., VanderHeide, J., LaCasse, T., Stanton, S., Coutermarsh, B., Barnaby, R., & Stanton, B. A. (2010). Arsenic inhibits SGK1 activation of CFTR Cl− channels in the gill of killifish, Fundulus heteroclitus. Aquatic Toxicology, 98(2), 157–164. 10.1016/j.aquatox.2010.02.001

75. Shaw, J. R., Hampton, T. H., King, B. L., Whitehead, A., Galvez, F., Gross, R. H., Keith, N., Notch, E., Jung, D., Glaholt, S. P., Chen, C. Y., Colbourne, J. K., & Stanton, B. A. (2014). Natural Selection Canalizes Expression Variation of Environmentally Induced Plasticity-Enabling Genes. Molecular Biology and Evolution, 31(11), 3002–3015. 10.1093/molbev/msu241

76. Soneson, C., Love, M. I., & Robinson, M. D. (2016). Differential analyses for RNA-seq: Transcript-level estimates improve gene-level inferences. F1000Research, 4, 1521. 10.12688/f1000research.7563.2

77. Sterner, R., Hagemeier, D., Smith, W., & Smith, R. (1993). Phytoplankton nutrient limitation and food quality for Daphnia. Limnology Oceanography, 38(4).

78. Sterner, R. W., & Elser, J. J. (2002). Ecological stoichiometry: The biology of elements from molecules to the biosphere. Princeton University Press.

79. Tam, L. M., Price, N. E., & Wang, Y. (2020). Molecular Mechanisms of Arsenic-Induced Disruption of DNA Repair. Chemical Research in Toxicology, 33(3), 709–726. 10.1021/acs.chemrestox.9b00464

80. Tam, L. M., & Wang, Y. (2020). Arsenic Exposure and Compromised Protein Quality Control. Chemical Research in Toxicology, 33, 1594–1604.

81. Tang, D., Chen, M., Huang, X., Zhang, G., Zeng, L., Zhang, G., & Wang, Y. (2023). SRplot: A free online platform for data visualization and graphing. PLoS One, 18(11), e0294236.

82. Theegala, C. S., Carriere, P. E., Francis, A., Tate, T., & Suleiman, A. A. (2006). Influence of primary producers on bioavailability of desorption resistant organic pollutants in the sediments. Soil and Sediment Con tamination: An International Journal, 15(3), 299–314. 10.1080/15320380600646373

83. Thompson, L. L., Rutherford, K. A., Lepage, C. C., & McManus, K. J. (2021). The SCF Complex Is Essential to Maintain Genome and Chromosome Stability. International Journal of Molecular Sciences, 22(16), 8544. 10.3390/ijms22168544

84. Twiss, M. R., & Nalewajko, C. (1992). INFLUENCE OF PHOSPHORUS NUTRITION ON COPPER TOXICITY TO THREE STRAINS OF *SCENEDESMUS ACUTUS* (CHLOROPHYCEAE)^1^. Journal of Phycology, 28(3), 291–298. 10.1111/j.0022-3646.1992.00291.x

85. Vahter, M.E., (2002). Mechanisms of arsenic biotransformation, Toxicology 181–182: 211-217.

86. Vahter, M. E. (2007). Interactions between arsenic-induced toxicity and nutrition in early life. Journal of Nutrition, 137(12), 2798–2804. 10.1093/jn/137.12.2798

87. Vahter, M. E., & Marafante, E. (1987). Effects of low dietary intake of me thionine, choline or proteins on the biotransformation of arsenite in the rabbit. Toxicology Letters, 37(1), 41–46.

88. Vandenbrouck, T., Soetaert, A., Van Der Ven, K., Blust, R., & De Coen, W. (2009). Nickel and binary metal mixture responses in Daphnia magna: Molecular fingerprints and (sub)organismal effects. Aquatic Toxicology, 92(1), 18–29. 10.1016/j.aquatox.2008.12.012

89. Villa-Bellosta, R., & Sorribas, V. (2010). Arsenate transport by sodium/ phosphate cotransporter type IIb. Toxicology and Applied Pharma cology, 247(1), 36–40. 10.1016/j.taap.2010.05.012

90. Villarroel, M. J., Sancho, E., Andreu-Moliner, E., & Ferrando, M. D. (2009). Biochemical stress response in tetradifon exposed Daphnia magna and its relationship to individual growth and reproduction. Science of the Total Environment, 407(21), 5537–5542.

91. Vighi, M., Altenburger, R., Arrhenius, Å., Backhaus, T., Bödeker, W., Blanck, H., Consolaro, F., Faust, M., Finizio, A., Froehner, K., Gramatica, P., Grimme, L. H., Grönvall, F., Hamer, V., Scholze, M., & Walter, H. (2003). Water quality objectives for mixtures of toxic chemicals: Problems and perspectives. Ecotoxicology and Environ mental Safety, 54(2), 139–150.

92. Walton, F. S., Waters, S. B., Jolley, S. L., LeCluyse, E. L., Thomas, D. J., & Styblo, M. (2003). Selenium compounds modulate the activity of recombinant rat AsIII-methyltransferase and the methylation of arsenite by rat and human hepatocytes. Chemical Research in Toxicology, 16(3), 261–265. 10.1021/tx025649r

93. Weider, L. J., Glenn, K. L., Kyle, M., & Elser, J. J. (2004). Associations among ribosomal (r)DNA intergenic spacer length variation, growth rate, and C:N:P stoichiometry in the genus Daphnia. Limnology and Oceanography, 49, 1417–1423. 10.4319/lo.2004.49.4_part_2.1417

94. Wingett, S. W., & Andrews, S. (2018). FastQ Screen: A tool for multi-genome mapping and quality control. F1000Research, 7, 1338. 10.12688/f1000research.15931.2

95. World Health Organization. (2022, December 7). *Arsenic*. https://www.who.int/news-room/fact-sheets/detail/arsenic

96. Xu, Z., Li, Y., Li, M., & Liu, H. (2021). Transcriptomic response of *Daphnia magna* to nitrogen- or phosphorus-limited diet. Ecology and Evolution, 11(16), 11009–11019. 10.1002/ece3.7889

97. Yang, C., & Frenkel, K. (2002). Arsenic-Mediated Cellular Signal Transduction, Transcription Factor Activation, and Abberant Gene Expression: Implications in Carcinogenesis. *Journal of Environmental Pathology*, Toxicology, and Oncology, 21(4), 331–342.

98. Yang, L., Kemadjou, J. R., Zinsmeister, C., Bauer, M., Legradi, J., Müller, F., Pankratz, M., Jäkel, J., & Strähle, U. (2007). Transcriptional profiling reveals barcode-like toxicogenomic responses in the zebrafish embryo. Genome Biology, 8(10), R227. 10.1186/gb-2007-8-10-r227

99. Yang, Q., & Fukamizo. (2019). Targeting Chitin-containing Organisms (Vol. 1142).

100. Zhang, H., Yang, L., Ling, J., Czajkowsky, D., Wang, J.-F., & Zhang, X.-W. (2015). Systematic identification of arsenic-binding proteins reveals that hexokinase-2 is inhibited by arsenic. PNAS, 112(49).

