## Supplementary figures and images for "Role of Nutritional Status on Arsenic Toxicity in *Daphnia pulex:* A Transcriptomic Perspective on Individual and Interactive Effects"

### Supplemental Figure 4

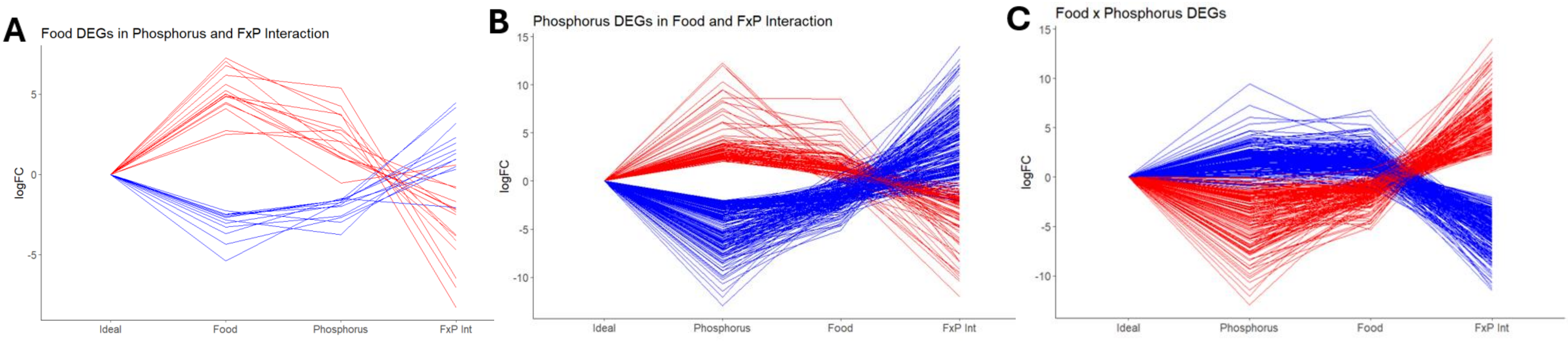
