## Supplemental Material for "Role of Nutritional Status on Arsenic Toxicity in *Daphnia pulex:* A Transcriptomic Perspective on Individual and Interactive Effects"

229 Main Street, Keene, New Hampshire 03435

| **Supplemental Table 1: Sequencing Statistics** | | | | | | |
| --- | --- | --- | --- | --- | --- | --- |
| **Sample** | **Group** | **Raw Reads** | **Surviving Reads** | **Survival Rate (%)** | **Mapped Reads** | **Mapping Rate (%)** |
| **Sample_1** | HP_3_As | 2,987,090 | 2,952,882 | 98.86 | 2,321,535 | 78.62 |
| **Sample_3** | HP_0.1_As | 2,208,441 | 2,178,504 | 98.64 | 1,602,862 | 73.58 |
| **Sample_8** | HP_3_0As | 3,433,153 | 3,398,243 | 98.98 | 2,164,933 | 63.71 |
| **Sample_9¹** | HP_3_0As | 38,397 | 27,093 | 70.56 | 8,285 | 30.58 |
| **Sample_14** | HP_0.1_0As | 2,806,413 | 2,777,703 | 98.98 | 1,946,469 | 70.07 |
| **Sample_20** | HP_0.1_As | 2,473,856 | 2,438,278 | 98.56 | 1,860,418 | 76.3 |
| **Sample_25** | LP_0.1_As | 1,502,513 | 1,475,558 | 98.21 | 1,165,076 | 78.96 |
| **Sample_28** | LP_3_As | 1,743,588 | 1,714,804 | 98.35 | 1,335,511 | 77.88 |
| **Sample_29** | HP_0.1_As | 2,444,216 | 2,411,713 | 98.67 | 1,594,281 | 66.1 |
| **Sample_31** | LP_3_0As | 1,642,452 | 1,613,426 | 98.23 | 1,003,051 | 62.17 |
| **Sample_33** | LP_3_As | 2,738,476 | 2,704,981 | 98.78 | 1,695,385 | 62.68 |
| **Sample_34** | LP_0.1_As | 3,390,260 | 3,348,961 | 98.78 | 1,843,206 | 55.04 |
| **Sample_35** | HP_3_As | 2,926,483 | 2,883,244 | 98.52 | 1,822,008 | 63.19 |
| **Sample_36** | HP_0.1_0As | 3,558,805 | 3,524,957 | 99.05 | 1,938,216 | 54.98 |
| **Sample_37** | LP_0.1_0As | 2,336,122 | 2,306,028 | 98.71 | 1,337,146 | 57.98 |
| **Sample_38** | HP_3_As | 2,265,250 | 2,224,555 | 98.2 | 1,343,546 | 60.39 |
| **Sample_43** | LP_0.1_0As | 1,988,726 | 1,958,119 | 98.46 | 1,207,904 | 61.69 |
| **Sample_46** | HP_0.1_0As | 430,896 | 417,879 | 96.98 | 200,189 | 47.91 |
| **Sample_48** | LP_3_As | 1,173,167 | 1,152,109 | 98.21 | 727,856 | 63.18 |
| **Sample_49** | LP_3_0As | 3,191,985 | 3,152,110 | 98.75 | 2,077,957 | 65.92 |
| **Sample_50** | LP_0.1_As | 914,575 | 897,585 | 98.14 | 538,182 | 59.96 |
| **Sample_51** | LP_0.1_0As | 1,528,332 | 1,505,237 | 98.49 | 712,321 | 47.32 |
| **Sample_52** | LP_3_0As | 621,549 | 603,447 | 97.09 | 391,485 | 64.87 |
| **Sample_57** | HP_3_0As | 2,455,887 | 2,423,464 | 98.68 | 1,779,878 | 73.44 |
| **Total** |  | **50,800,632** | **50,090,880** |  | **32,617,700** |  |
| **Overall %** |  |  |  | **98.60%** |  | **65.12%^[1]^** |

**Table S1: Sequencing Statistics**

This table includes raw and processed sequencing information and statistics for this study. Group descriptions: *PhosphorusQuality(HP or LP)_FoodQuantity(3 or 0.1)_Arsenic(As or 0As).* Surviving Reads provides the total number of reads that survived trimming. Survival rate calculated as surviving reads/raw reads x 100. Mapped reads provides the total number of reads that mapped to the reference transcriptomic index. Mapping rate calculated as mapped reads/raw reads x 100.

**Fig. S1: Low log transformed normalized counts indicate lack of normalization for Sample 9**

Figure S1 shows the Trimmed Mean of M-values (TMM) normalization method from *EdgeR* to calculate counts per million (CPM) for each sample.
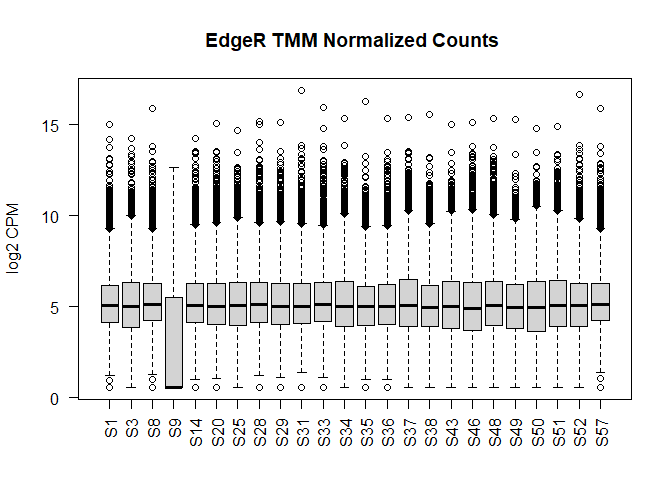

Figure S1: Bar plot exhibiting the median log_2_CPM values for each sample initially included in this study. All samples except S9 were able to be normalized using this method as well as other samples exhibiting low CPMs (see Table S1 for samples 46 and 51).

**Fig. S2: Hierarchical clustering reveals Sample 9 is an outlier**

Samples were clustered based on Euclidean distances calculated from normalized counts for each sample. The dendrogram shows 8 major clades where 6 are separately by two replicates of the same treatment (S35 & S38, S37 & S51, S31 & S52, S8 & S57, S28 & S33, S20 & S29). One treatment (HP_0.1_0As: S14, S36, S46) did not cluster together. Two of the clades (S43 & S50 and S3 & S25) are replicates from different treatments. Sample 9 separates from all samples.

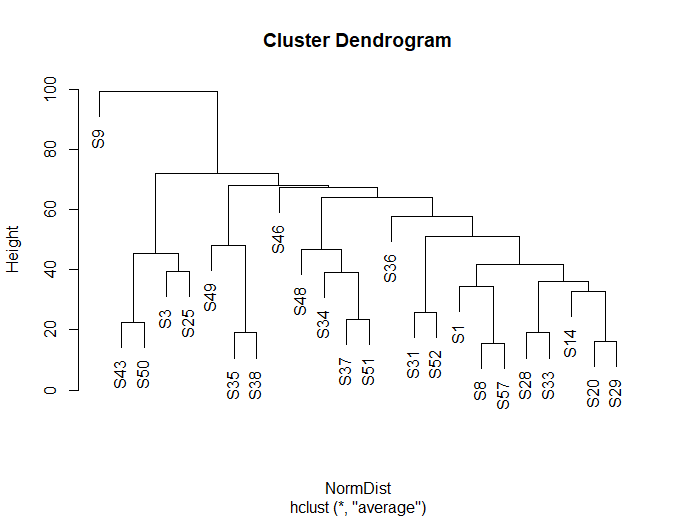

Figure S2: Hierarchal clustering dendrogram of each sample. Sample 9 (S9) separated from all samples prompting its removal from downstream analysis. Lack of clustering was indicated within the HP_0.1_0As samples: S14, S36, and S46.

**Fig. S3 Principal Component Analysis (PCA) of RNA-sequencing samples**

Fig. S3 PCA plot of *D. pulex* biological RNA-sequencing replicates. The shapes indicate the presence/absence of arsenic while the color indicates the combination of low phosphorus and low food quantity treatment. The first principal component indicates that 26.3% separate based on phosphorus rather than arsenic or food.

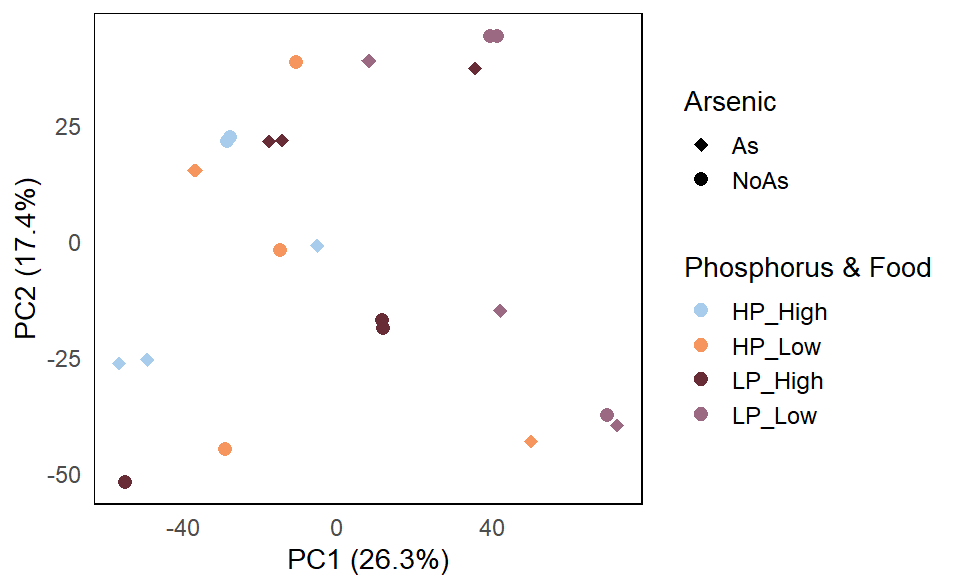

Figure S3: Principal component analysis (PCA) of D. pulex treatment replicates. Due to the nature of design, the low phosphorus and low food quantity treatments were combined for ease of interpretation. Shape indicates whether arsenic was present (As, diamond) or absent (NoAs, circle).The combination of phosphorus and food are dictated by color: LightBlue (HP_High), Orange (HP_Low), DarkRed (LP_High), and Purple (LP_Low).

**Table S2: Table Exhibiting Total and Annotated Number of Differentially Expressed (DE) Genes from the Generalized Linear Model**

Table S2 showcases the number of DE genes for each main effect and each binary interaction from the full generalized linear model (glm): *~ Food + Phos + Arsenic + Food*Phos + Food*Arsenic + Phos*Arsenic + Food*Arsenic*Phos*. This analysis focused on the interpretation of binary interactions. The first columns exhibits the total number of DE genes from the glm. Annotations for protein coding genes revealed that nearly all main effects and binary interactions had several lacking annotations due to the presence of long non-coding RNAs (lcnRNA) and confirmed by NCBI blast prior to removal. The second column lists the number of DE genes after lncRNA removal.

| **Main Effect or Interaction Term** | **Total Number of DE Genes** | **Number of Annotated DE Genes after lncRNA removal** |
| --- | --- | --- |
| **Arsenic** | 51 | 46 |
| **Phosphorus** | 321 | 295 |
| **Food** | 27 | 27 |
| **Phosphorus x Arsenic** | 456 | 435 |
| **Food x Arsenic** | 2 | 2 |
| **Food x Phosphorus** | 428 | 408 |

**Tables S3– S8: Annotation Files for each Main Effect and Binary Interaction Term Differentially Expressed Genes from the generalized linear model.**

Tables S3 – S8 contain annotation files for each main effect and binary interaction term set of differentially expressed gene set interpreted in this study. Annotations are from the National Center for Biotechnology Information (NCBI). The order of annotated differential expression begins with main effects arsenic (Table S3), phosphorus (Table S4), food (Table S5) and followed by binary interactions phosphorus x arsenic (Table S6), food x arsenic (Table S7), and food x phosphorus (Table S8). Each table contains the following columns:

- **Genes**: unique NCBI gene symbol identifier with the prefix “LOC”
- **logFC**: abbreviation for log_2_fold change of gene expression after treatment
- **PValue**: raw p-value from appyling likelihood ratio tests
- **FDR**: adjusted p-value after applying multiple test correction with Benjamini-Hochberg (BH) factor
- **DGE**: categorical designation based on logFC > 2and FDR < 0.05
- **Group**: categorical designation based on main effect or binary interaction status (arsenic, phosphorus, food, etc.)
- **Protein.Name**: NCBI annotation for a corresponding gene
- **GeneID**: unique NCBI numerical identifier for each gene

**Table S9: Table of Collective Pathway Activation Analysis Results for Main Effects**

Table S9 exhibits the pathway activation results for arsenic, food, and phosphorus main effects. Each main effect is sorted by lowest FDR. The table includes the following columns:

- **Total Genes**: total number of genes detected as significant for a specific KEGG pathway and meeting the minimum of 4 genes for a significant pathway
- **Activated Genes**: categorized by a positive median logFC > 1 and FDR < 0.05
- **Repressed Genes**: categorized by a negative median logFC < -1 and FDR < 0.05
- **Percentage of Genes Activated:** calculating the activation of significant pathway via the number of activated genes/ total genes x 100.
- **Median LogFC**: the median of the collective logFC from all genes determined to be significant based on that pathway
- **P-value**: raw p-value from applying the binomial statistical test
- **FDR**: adjusted p-value after applying multiple testing correction with Benjamini-Hochberg (BH) factor

**Fig. S4 Low Food x Low Phosphorus Gene Expression Response**

Figure S4 exhibits that the DE gene sets for (A) low food, (B) low phosphorus, and (C) the low food x low phosphorus interaction also produce an antagonistic response. As the focus of the paper was arsenic-nutrient interactions, the dual nutrient interaction was placed in the supplemental materials. This multipaneled figure has been uploaded separately as a pdf. This caption serves as a chronological placeholder for where this figure is referenced in the manuscript.

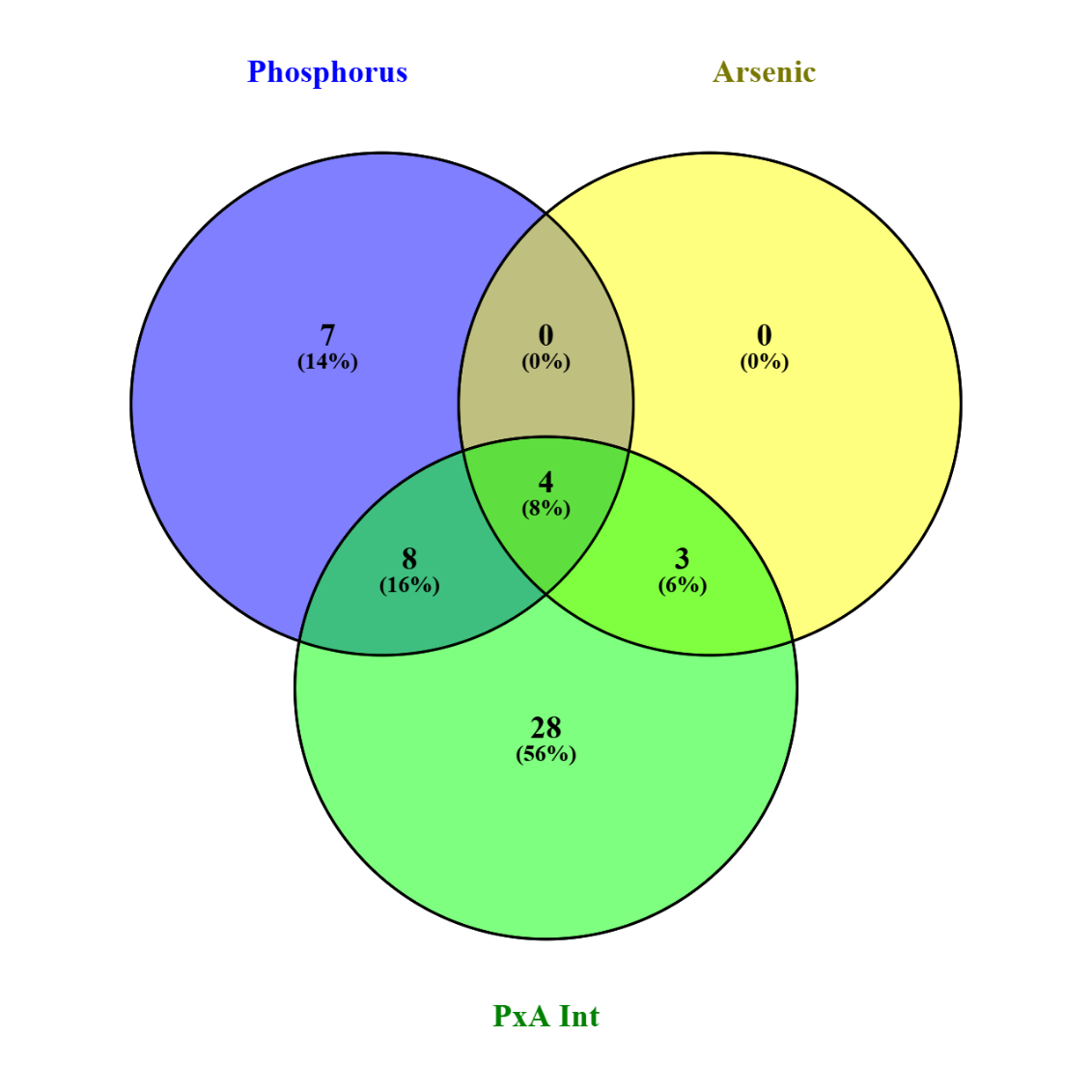
**Fig. S5 Venn Diagram of Overlapping in Significant Pathways Between Main Effects Low Phosphorus, Arsenic, and the Low Phosphorus x Arsenic Interaction**

Figure S5: Assessment in the number of significant pathways determined by Pathway Activation Analysis among two main effects, low phosphorus and arsenic, and their subsequent interaction term (phosphorus x arsenic) reveal increased overlap in shared pathways. Low Phosphorus (19 total sig. pathways) shares a total of 12 with Arsenic resulting in only 7 pathways being unique to this main effect. All of the significant pathways for the arsenic main effect (7 total sig. pathways) were shared with low phosphorus and the subsequent interaction. Low phosphorus x arsenic interaction indicates that 65% of the sig. pathways (28/43) are unique to the interaction and collectively share 15 pathways across both low phosphorus and arsenic.

1. Sample_9 was excluded from downstream analysis due to low mapping rates and contamination. According to NCBI Blast, overrepresented sequences matched *Daphnia pulex* mitochondrial DNA (e.g. GCCCCAACAAAATTTCATCATAATTTTTGTTATAGAAAGTAACCTAAAAC) and *Ceriodaphnia* sp. ribosomal RNA (e.g. GCACCTTGCTAATTTCTTAATCCAACATCGAGGTCGCAAACCTTTTTATC).
